# Ensembles of *in silico* structures enable T cell peptide-MHC binding prediction

**DOI:** 10.64898/2026.07.14.738340

**Authors:** O Lyudovyk, JA Levine, M Pathil, S Martis, A Streltsov, Z Sethna, Y Elhanati, VP Balachandran, Q Morris, BD Greenbaum

## Abstract

Adaptive immunity relies on T-cell receptor (TCR) recognition of peptides presented by the major histocompatibility complex (pMHC). Accurate prediction of TCR:pMHC binding pairs from sequence data remains a longstanding challenge in computational immunology, limiting the development of precision immunotherapies like cancer vaccines and adoptive cell therapies. Here, we present enFoldX (**en**semble of **Fold**ed comple**X**es), a structure-based approach leveraging biophysical characterization of AlphaFold3-generated ensembles to classify TCR:pMHC sequence pairs as cognate versus non-cognate. Unlike previous methods reliant on only sequence data or a single, static predicted structure, enFoldX extracts features from an entire generated ensemble with a custom focus on the biophysical binding interface. Our model distinguishes T cell reactivity between peptides differing by a single amino acid substitution, the resolution required for cancer neoantigens, and generalizes to unseen peptides, MHCs, and TCRs, a major objective for artificial intelligence (AI) in immunology. Our performance on these crucial tasks demonstrates that diverse, structural sampling of biophysical interactions over an ensemble is fundamental for accurate AI-driven binding predictions and offers lessons for efficient future data generation to improve models. Our findings therefore offer a scalable framework to accelerate therapeutic binder design, and we provide access to a publicly available code repository.

## INTRODUCTION

T cells are central to immunity, recognizing and killing virus infected cells, and playing a critical role in immune surveillance of tumors and their clearance by immunotherapies^1^. Their specificity is encoded by T-cell receptors (TCRs) which can recognize epitopes, linear peptides presented on the cell surface by major histocompatibility complex (MHC) molecules. Although V(D)J recombination generates 10^9^-10^10^ distinct TCR sequences in a typical human^2,3^, only tens to thousands of TCRs are estimated to recognize any given epitope. While the adaptive immune system evolved to defend against “nonself” pathogens^4^, T cells can also recognize cancer neoantigens, peptides derived from mutations in a tumor cell, and viral antigens that differ from self-peptides by only one amino acid^5–8^.

High-throughput sequencing of TCRs has fueled a growing body of computational methods for predicting TCR:pMHC binding pairs from sequence data^9,10^, spanning graph-based models, large language models (LLMs), and AI-based structural prediction and docking methods^11–15^. While some have shown promise, generalization to novel pairings remains a fundamental challenge with no clear consensus solution^16,17^. Predicting which TCRs bind a given peptide-MHC (pMHC) complex from sequence data has broad applications, from understanding viral clearance and self-tolerance, to identifying biomarkers for autoimmune diseases, and designing personalized cancer vaccines and T cell therapies. Doing so from sequence data alone has become a significant unsolved problem for AI in biology.

To predict TCR:pMHC pairing, suggestive results have come from methods leveraging protein structure prediction models. While these methods outperformed other approaches on unseen epitopes in a recent competition (IMMREP25)^18^, they rely on a single, high-confidence structure for each TCR:pMHC to make binding predictions. By doing so, these methods do not make full use of the ability of state-of-the-art structure prediction models to generate multiple predictions for a single input. This sampled ensemble, as we illustrate below, shows informative variability; relying on a single structure ignores this information and introduces high levels of uncertainty in binding predictions.

To address these issues, we developed enFoldX, which uses customized ensembles of predicted structures to assess binding. enFoldX extracts distributions of AlphaFold3^19^ (AF3) confidence features, including customized biophysical features, across many stochastically generated structures, and uses these distributions to train a binding classifier, analogous to an energy function in statistical physics. We provide access to a publicly available repository for running enFoldX, the first ensemble-based solution, to accelerate development, including a tutorial for predicting TCR specificity using feature ensembles.

We show that distributions of these customized features vary significantly between cognate and non-cognate TCR:pMHC pairs, as does the propensity to hallucinate physiologically implausible structures. Binary classifiers trained on ensemble distributions of features perform significantly better than both single-structure co-folding methods and sequence-based models on multiple independent datasets of TCR specificity. Importantly, enFoldX generalizes in a dataset agnostic way: models trained on human data generalize to other human datasets and even mouse TCR:pMHC pairs. In contrast, we show that sequence models only generalize to TCRs and epitopes that are highly similar to those in their training sets. Moreover, our structure-guided models trained on publicly available data can discriminate between unseen epitopes that are only one mutation away from self-peptides, a capacity critical to predicting mutation-derived cancer neoantigens for cancer vaccines and other precision immunotherapeutic applications^20–23^. We therefore conclude that biophysically customized, parallelized, structure-ensemble based approaches can unlock prediction of TCR:pMHC pairing, while holding general lessons for computational prediction of complex binding problems.

## RESULTS

### Structural ensembles of physically relevant features improve TCR:pMHC classification

enFoldX leverages AF3 to generate an ensemble of structural predictions and corresponding confidence estimates for TCR:pMHC complexes modeled with four subunits: peptide, MHCα chain, and TCR α and β chains (**Figure 1a**). For each complex, we generate a diverse ensemble of AF3-predicted structures using multiple seeds and multiple samples per seed. AF3, like other diffusion-based co-folding models, often generates plausible, false-positive structures for non-cognate TCR:pMHCs that are nearly indistinguishable from cognate structural predictions, particularly when the input sequence differ subtly (**Figure 1b**). However, our customized features, based on AF3 output, and ensembled over a diverse multitude of structures for each TCR:pMHC, enable discrimination between cognate and non-cognate pairs, even those that differ by a single mutation in the peptide (**Figure 1c**).

**Fig. 1.**
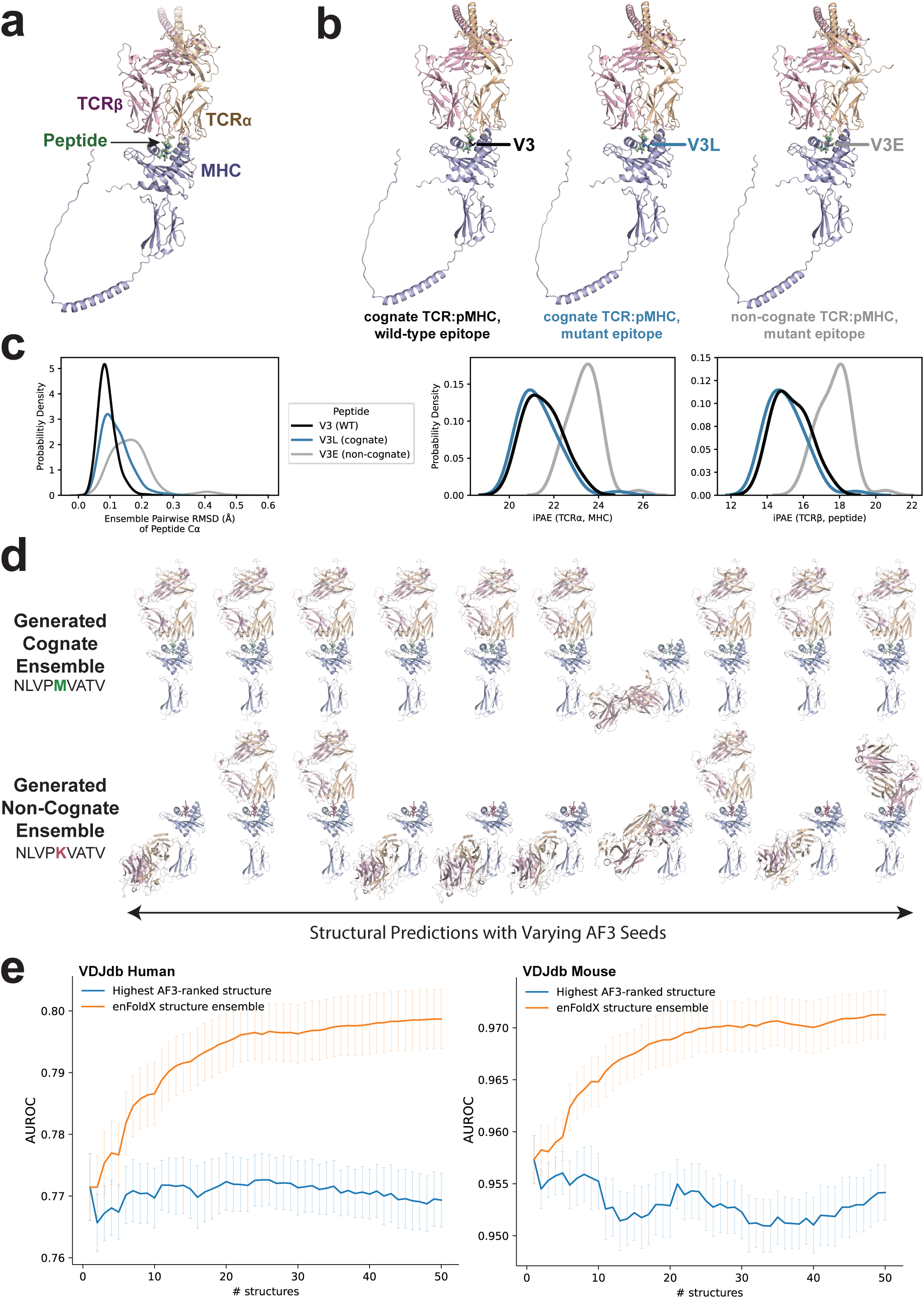
Ensembles of structures improve TCR specificity predictions. **a,** Representative AF3-predicted structure of TCR:pMHC complexes with four subunits: peptide, MHC α chain, and TCR α and β chains. **b,** AF3-generated plausible structures that are nearly indistinguishable for a TCR in complex with cognate (V3, V3L) and non-cognate (V3E) peptides presented on HLA-A*02:01. **c,** Probabilities of indicated ensemble output metrics for the same TCR:pMHCs shown in **b.** Ensembles for the non-cognate peptide (gray) exhibit higher levels of structural diversity measured by ensemble pairwise RMSD (**left**) and right-shifted interfacial iPAE distributions (**right**) relative to cognate peptides (black, blue). **d,** Representative structural ensembles for a TCR with a cognate pMHC (**top**) and a non-cognate pMHC (**bottom**) for a single input TCR:pMHC with varying seeds (also see **Extended Data Figure 1**). **e,** Comparison of performance (as measured by AUROC) on predicting cognate versus synthetically-generated non-cognate peptides for TCR binding in VDJdb (human and mouse) as a function of the number of structures used in the ensemble for enFoldX predictions (orange) and for a single structure with the highest AF3 score, which is the current paradigm in structure-based predictive models (blue). Error bars indicate standard error of the mean (SEM) across 5 cross validation folds. Data for **b-d** was sourced from TCRs specific to Human Cytomegalovirus wild-type NLV peptide (“NLVPMVATV”). Data for **b-c** is from TCR1 in the cross-reactivity dataset and data for **d** is from TCR 5-2 in the PTEAM dataset.

With each generated structure, the default AF3 pipeline outputs global and local confidence metrics including predicted alignment error (PAE), predicted local distance difference test (pLDDT), predicted template modeling (pTM) score, and contact probabilities. We directly extract AF3 confidence measures for the overall TCR:pMHC complex, each chain individually, and, when applicable, each pair of chains. To classify TCR:pMHC pairs as cognate or non-cognate, we engineered features to capture structurally relevant information, particularly at the TCR:pMHC binding interface. Previous work in *de novo* protein design has demonstrated the utility of interaction PAE (iPAE), or averages of PAE scores for pairs of inter-chain residues, in predicting protein-protein binding^24–26^. Building on this work, for some of enFoldX’s features, we defined the canonical TCR:pMHC interface to include iPAEs between all pairwise groups of chains as well as subchain iPAEs between the peptide and TCR complementarity-determining region 3 (CDR3) loops, the primary determinants of pMHC recognition (**Extended Data Figure 1a**).

To illustrate the advantages of structural ensembles, we analyzed the predictions of a TCR known to recognize a wild-type human cytomegalovirus (CMV)-derived epitope (“NLVP**M**VATV”) but not its mutant counterpart (“NLVP**K**VATV”, labelled as M5K)^27^. Of note, the ensembles for both the cognate and non-cognate TCR:pMHCs include a structural hallucination in which the TCR is misdocked in the position canonically occupied by β2-microglobulin in physiological TCR:pMHC binding (**Figure 1d, Extended Data Figure 1b**). However, the rate of this hallucination is far greater for the non-cognate ensemble in which seven out of ten randomly tested seeds exhibit the misdocked structure with high AF3-estimated error, demonstrating one reason why ensembles are needed for discrimination (**Extended Data Figure 1c**). Moreover, due to the previously mentioned high false positive rate for co-folding models, there are also many high-confidence structures in the non-cognate ensemble with low predicted error that, unlike hallucinations, resemble typical TCR:pMHC geometry with the TCR diagonally docked across the pMHC. Based on these findings, while a model only considering the high-confidence, plausibly docked M5K structure could misclassify the non-cognate pair, our ensemble-based approach captures a systematic difference in diversity across cognate and non-cognate structures, even when the only difference in input sequences is a single amino acid substitution in the peptide sequence.

In initial evaluations of our method, we chose the widely used benchmarking dataset VDJdb, the largest curated database of TCR sequences with known antigen specificity^28^. For many enFoldX-extracted features, we observed statistically significant differences between cognate and non-cognate pairs (**Extended Data Figure 1d**), suggesting that simple classifiers trained on this feature space can predict TCR specificity. We compared classification performance with our ensemble-based approach against the current paradigm of representing a given TCR:pMHC with a single structure for both human and mouse VDJdb data. Across both datasets, while performance saturates when only using the top-ranked structure with highest AF3 confidence (**Figure 1d, blue**), enFoldX performance consistently outperforms the single structure-based model and increases with additional structures included in the ensemble (**Figure 1d, orange**). We conclude that for challenging systems with flexible protein-protein interfaces like the TCR:pMHC complex, the integration of multiple stochastically generated samples is required to robustly predict binding.

### enFoldX architecture

enFoldX consists of four distinct steps run sequentially in a highly parallelized manner to classify a given TCR:pMHC as cognate or non-cognate (**Figure 2**). Using inputs of peptide, MHCα, and TCRα/β sequences, we first perform the default AF3 homology search to generate multiple sequence alignments (MSA) and select structural templates. We then run the AF3 diffusion-based inference pipeline to generate an ensemble of structural predictions and corresponding confidence estimates. Next, we extract structurally relevant confidence features, including those focused on the TCR:pMHC binding interface, from each predicted structure and generate a feature vector by computing the mean and standard deviation of each feature for all sampled structures in the ensemble. We applied this pipeline to generate a full feature set for the human VDJdb dataset and to train a classification model to predict cognate from non-cognate TCR:pMHC pairs. Parallelization of each step in our pipeline rendered large-scale TCR:pMHC classification tractable, speeding up predictions by a factor proportional to the amount of available compute (**Extended Data Figure 2**) with a trade-off between performance boost and computational cost of additional seeds.

**Fig. 2.**
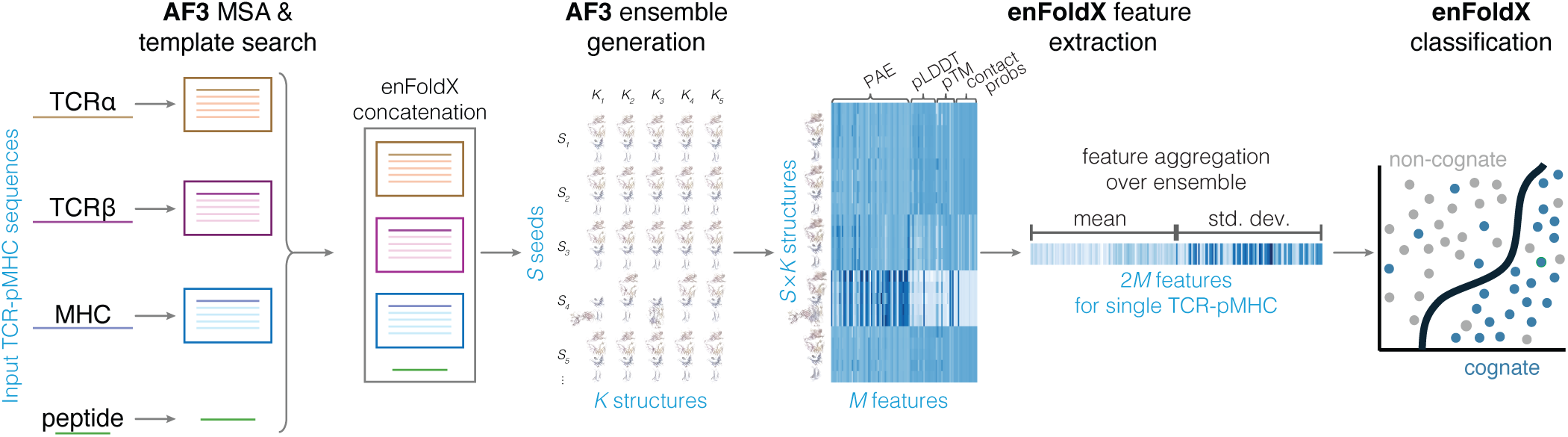
enFoldX architecture for structure ensemble based TCR:pMHC specificity prediction. Schematic of enFoldX ensemble-based prediction of TCR:pMHC binding. Input is a single sequence of each TCRα/β, MHCα, and peptide. The pipeline begins with AF3 MSA and template search followed by concatenation of MSA/template results, AF3 diffusion-based generation of structural ensembles, and feature extraction with aggregation over the ensemble. For a single input, the pipeline creates an ensemble of size *S* times *K*, where *S* is the number of input random seeds and *K* is the number of samples per seed. We extract *M*=106 features per structure and then summarize the feature distributions over ensemble using the mean and standard deviation of the feature values across all structures, resulting in 2*M* features per input (**Methods**). This feature set can then be used as inputs for fitting a classification model to labeled training data and evaluating it on unseen/held-out data.

We validated enFoldX on a broad range of prediction tasks and benchmarked performance against published, state-of-the-art (SOTA) TCR:pMHC prediction methods. We first assessed tasks for which published SOTA models perform well, largely focusing on the VDJdb datasets. We then evaluate enFoldX on two ongoing challenges for SOTA models: (i) generalization to new peptides, MHCs, and TCRs that substantially differ from those in the training set, and (ii) discrimination between TCR:pMHC pairs with single substitutions in the epitope.

### Biophysically customized features enable generalizability

As VDJdb consists exclusively of cognate TCR:pMHCs (positive examples), training our classification model required synthetic generation of non-cognate pairs (negative examples). To do so, we grouped TCR:pMHCs by MHC allele and swapped epitopes with “decoy” peptides identified by computational pMHC binding predictors (NetMHC4.0^29^ for mouse data and NetMHCPan4.1^30^ for human data) to be presented on the same MHC allele. This approach attempts to generate new TCR:pMHC pairs that are non-cognate due to inhibited TCR recognition of the pMHC, rather than failed MHC presentation of the peptide. Our non-cognate labeling of these synthetic decoys is based on the fact that a randomly paired TCR:pMHC is statistically unlikely to bind.

Combining our decoy non-cognate pairs with cognate TCR:pMHCs from VDJdb, we compared enFoldX classification performances across human and mouse datasets (**Figure 3**). We performed ten-fold cross-validation by TCR and epitope with a L2-regularized logistic regression classification model (Ridge Regression)^31^, splitting folds by unique TCR or epitope sequences to mitigate data leakage between training and test data. For a classifier trained on human data (**Figure 3a, left**), we achieved AUCs, averaged across ten test folds of 0.82 for TCR-wise cross-validation and 0.73 for epitope-wise cross-validation. We observed similar trends for a separate classifier trained on mouse data (**Figure 3a, center**) with mean AUCs of 0.98 and 0.95 by TCR and epitope, respectively. Our stronger performance on TCR-wise cross-validation for both human and mouse classifiers indicates that training examples of TCR cross-reactivity, in which a single TCR recognizes multiple pMHC antigens, may confer more predictive power than examples of TCR degeneracy, in which a single pMHC is recognized by multiple TCRs. Additionally, our model trained exclusively on human data can classify TCR:pMHCs from mouse data (**Figure 3a, right**), demonstrating transfer learning across species for the first time.

**Fig. 3.**
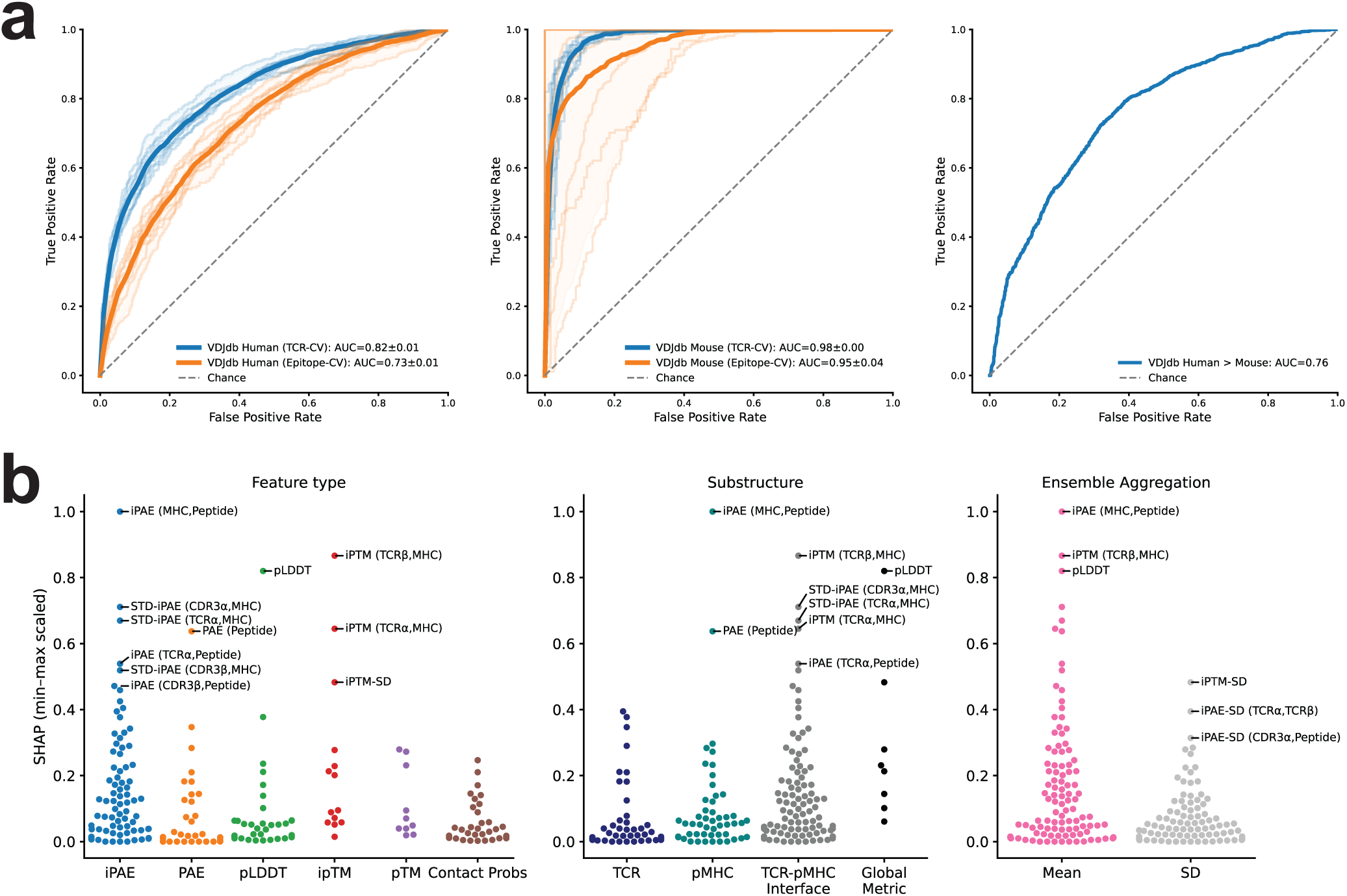
enFoldX generalizes across novel TCRs and epitopes in VDJdb, within and across species. **a,** ROC curves for classification of cognate (positive) vs. synthetic non-cognate (decoy negative) TCR:pMHCs in human (**left**) and mouse (**center**) VDJdb datasets and for cross-species generalization (**right**). For individual human and mouse datasets, AUROC values are shown as mean and confidence intervals over folds using ten-fold cross-validation, splitting folds based on unique TCR (blue) or epitope (orange) sequences. For cross-species generalization, performance is shown for classification of TCR:pMHCs in mouse VDJdb using an enFoldX model trained exclusively on human VDJdb data. Results are shown for L2-regularized Logistic Regression classification models. **b,** Normalized feature importances by SHAP (**Methods**) for the human VDJdb classification task shown on the left in **a**. Features are stratified by feature type (**left)**, by substructure (**center)**, or by ensemble summary statistic (**right**).

To understand which metrics are most informative for our human classifier, we computed Shapley Additive exPlanations (SHAP)^32^ values to quantify the average marginal contribution of each feature (**Figure 3b**). Metrics focused on the TCR:pMHC and pMHC binding interfaces appeared to be crucial for classification with the dominant signal from iPAEs and substantial contributions from iPTMs. Interestingly, the most informative global metric was our pLDDT feature, a summary of the local confidences in the overall complex for a structural ensemble. We also note that both means and standard deviations contributed predictive value, indicating that the distributions of features across ensembles carry important, non-redundant information. Collectively, this analysis reaffirms that our customized emphasis on points of biophysical contact at the TCR:pMHC interface, our diverse feature set integrating local and global confidence measures, and our aggregation strategy across predicted ensembles are all needed to enable accurate classification.

As our decoy method relies on netMHC-based pMHC binding predictors, we explored a simpler, “permuted” approach in which we group TCR:pMHCs by MHC allele and randomly permute TCRs and epitopes within each group. This ensures that the permuted non-cognates have experimentally confirmed MHC presentation so functional differences between TCR:pMHCs can only be attributed to altered TCR recognition. Like the decoy method, randomly paired TCR:pMHCs are assumed to be non-cognate. Repeating the experiments shown in **Figure 3** for our permuted approach, we observed the same overall trends regarding higher AUCs for cross-validation by TCR versus epitope and generalizability from our human classifier to mouse data (**Extended Data Figure 3a**). However, across all classification tasks, our permuted performance is lower than that of the decoy method. Feature importance analysis revealed that while interfacial features like iPAEs and iPTMs are still the most informative measures, the permuted classifier relies more heavily on features specific to the TCR:pMHC interface, such as interactions between the peptide and CDR3β or peptide and TCRα (**Extended Data Figure 3b**). Based on these findings, we posit that our decoy model, which emphasizes enFoldX features for both the pMHC and TCR:pMHC interfaces, better predicts MHC presentation and TCR recognition conditioned on MHC presentation. This enables better generalization to datasets with unseen TCRs and epitopes, as in the human to mouse classification task, and peptide mutational scans, as we later describe, where a loss in TCR activity may be due to failed MHC presentation.

### Generalization to unseen data and peptide mutational scans

We next evaluated enFoldX performance on two ongoing challenges for the field: (i) generalization to out-of-distribution TCRs or pMHCs and (ii) TCR discrimination between epitopes that only differ by a single substitution. To do so, we selected a broad range of datasets varying in size, sequence diversity, class balance, and MHC allele representation (**Extended Data Figure 4, Methods**). Across datasets, our most informative features, including custom chain and subchain iPAEs describing CDR3-peptide interactions, differed between cognate and non-cognate TCR:pMHC pairs, indicating enFoldX could predict TCR specificity across these different validation sets (**Extended Data Figure 5**).

We first assessed enFoldX on unseen data from a recent VDJdb release in September 2025 (“VDJdb 09/2025”). Our model, trained on data released in February 2025, demonstrated strong performance with an AUC of 0.86 on new TCRs and 0.77 on new epitopes (**Figure 4a**). Given recent suggestions that several TCR:pMHC pairs labeled positive in VDJdb may be unreactive^33^, we also evaluated enFoldX on TCRvdb, a smaller functionally validated TCR library. Trained on a subset of the February and September 2025 VDJdb releases to exclude any TCRvdb records, the model scored an AUC of 0.75 for distinguishing experimentally confirmed cognate pairs from those that failed functional validation. The ability of enFoldX to predict for other datasets while trained on the full VDJdb, which contains false positives^33^, suggests that our diverse feature set capturing structural information over ensembles can identify meaningful signal, even when the training set contains false positives (**Figure 4a**).

**Fig. 4.**
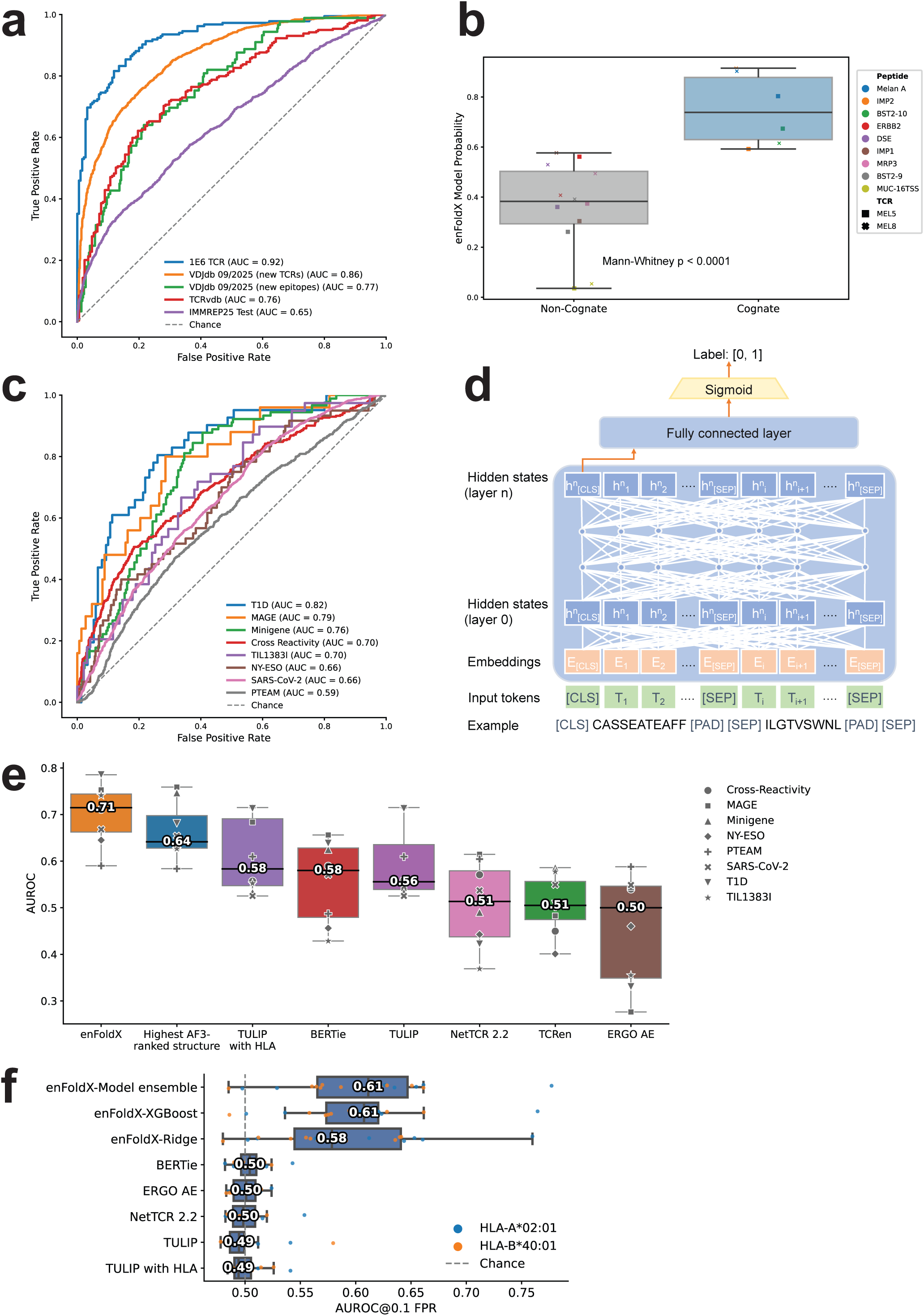
enFoldX demonstrates consistently high performance across experimental TCR specificity datasets and peptide mutational scans. **a,** enFoldX-Ridge model performance on the cognate epitopes for the hyper cross-reactive 1E6 TCR (blue), VDJdb updated 09/2025 (orange, green), experimental data from TCRvdb (red), and the evaluation set from the IMMREP25 competition (purple). **b,** enFoldX-Ridge model predictions for reactivity of MEL5 and MEL8 TCRs to three cognate epitopes (Melan A, IMP2, and BST2-10) and several non-cognate epitopes (Mann-Whitney U test, p<0.0001). **c,** enFoldX-Ridge model performance on eight independent peptide mutational scan datasets. **d,** Architecture and input example for a sequence-based model, BERTie. **e,** Performance comparison of enFoldX-Ridge and indicated methods on the mutational scan datasets from **c.** Highest AF3-ranked structure model (in blue) is trained using the 106 confidence and contact probability features as described but extracted from only the top-ranked predicted AF3 structure for each TCR:pMHC complex. The horizontal line on the boxplots represents the median of AUCs for all mutational scan datasets. **f,** Performance comparison of enFoldX models and indicated methods on the IMMREP25 dataset. Boxplots show the distribution of AUROC@0.1 FPR – the area under the ROC curve up to FPR = 0.1 – across independent epitopes, with median values indicated.

It is important to note that both VDJdb releases include examples of TCR degeneracy, with some epitopes recognized by many TCRs, but limited examples of TCR cross-reactivity with a single TCR recognizing multiple antigens. As accurately scoring promiscuous TCRs is important for TCR:pMHC classification^34^, we tested enFoldX on 1E6, a well-characterized TCR estimated to bind to over one million peptides^35^, as well as on MEL5 and MEL8, multireactive TCRs that recognize multiple tumor-associated antigens^36^. For 1E6, we assessed performance on a dataset of 185 known cognate epitopes with synthetically generated non-cognate pairs, scoring an AUC of 0.92 (**Figure 4a, Methods**). For the MEL TCRs, we found a significant difference (p<0.0001) in enFoldX-predicted binding probabilities between experimentally validated cognate and non-cognate examples (**Figure 4b**). Taken together, these results indicate that although enFoldX training data was limited in the representation of cross-reactivity, the model can correctly classify TCR:pMHCs with promiscuous TCRs.

We then evaluated enFoldX on newly released data from IMMREP25^18,37^, a community-organized challenge for predicting TCR specificity. Focusing on generalizability to unseen antigens, the original IMMREP25 dataset includes data for 1,000 TCRs with 20 unseen peptides with computationally predicted presentation by one of two common human leukocyte antigen (HLA) alleles (HLA-A*02:01 or HLA-B*40:01). All included epitopes differ from published peptides by a Levenshtein distance of at least four amino acids. We note that while the original dataset with 10,000 records was released, the precise composition of the IMMREP25 evaluation set, approximately 75% of the original data, was not disclosed and therefore cannot be reconstructed. For this reason, we report results here for all records included for the 18 pMHC antigens in the IMMREP25 private leaderboard (**Methods**). We also note that both MHC presentation and TCRα/β-peptide pairings in IMMREP25 were computationally inferred rather than experimentally determined, so it almost certainly contains some non-cognate TCR:pMHCs mislabeled as cognate. Nonetheless, we first evaluated enFoldX performance with a ridge regression classifier trained on all human examples from the February and September 2025 VDJdb releases with both decoy and permuted negatives, scoring an overall AUROC of 0.65 (**Figure 4a**). A model ensemble consisting of a uniformly weighted combination of Ridge Regression^31^, XGBoost^38^, and Random Forest^39^ classifiers achieved a macro-AUROC of 0.66 and median AUROC with a maximal false positive rate of 10% (“AUROC@0.1 FPR”) of 0.61, outperforming all sequence- and structure-based methods reported in IMMREP25 (**Figure 4f, Extended Data Figure 8a**).

Lastly, we tested enFoldX performance on distinguishing cognate from non-cognate TCR:pMHCs that differ only by single amino acid substitutions in the epitope — a critical task in the development of precision immunotherapies. We surveyed eight mutational scan datasets (**Methods**), each comprising a TCR:pMHC library that maps how peptide point mutations in a validated cognate TCR:pMHC impact its binding or T cell activation levels. Though these datasets vary by experimental design, most assays were conducted at single peptide concentrations rather than titrations and reported half-maximal effective concentration (EC_50_) values, or a similar assay-dependent measurement. As a single concentration is insufficient to rigorously measure EC_50_ values, and the measurements are specific to the assay experimental details, we note there is a strong likelihood of noise in assay comparisons, since the place of the assay along an EC50 binding curve cannot be established by a single concentration assay. Nevertheless, enFoldX performance averaged to a macro-AUC of 0.71 (**Figure 4c, Methods**), demonstrating its sensitivity to the impact of subtle peptide mutations on TCR recognition. Consistent with the wide variations in experimental design, our performance varied widely with AUCs ranging from 0.59 to 0.82.

Taking a closer look at enFoldX performance on this specific task, we performed additional experiments with our previously published mutational scan dataset^21^ that, unlike the other datasets, measured TCR activation at several peptide concentrations for seven TCRs cognate to three known pMHC antigens (**Extended Data Figure 6a**). We applied a cross-validation strategy where we, in turn, withheld one of the nine mutated positions or one of the seven TCRs. Here, enFoldX performs significantly better than random at classifying TCR:pMHCs with single amino acid substitutions for six of nine mutated positions and four of seven TCRs (**Extended Data Figure 6b**, statistical significance was assessed using a one-sided permutation test with 10,000 random label permutations and Phipson–Smyth-corrected p-values). Notably, all seven TCRs and all nine mutated positions achieved AUCs greater than 0.5. Regarding model architecture, we evaluated several machine learning classifiers, including XGBoost and Random Forest, finding best generalizability from a VDJdb-trained model to our cross-reactivity data using Ridge Regression (**Extended Data Figure 6c**). To understand which feature subsets are most informative for generalization to cross-reactivity, we also considered a family of models trained on subsets of our feature space, comparing various AF3-derived confidence features (**Extended Data Figure 6d**) and feature average versus standard deviation metrics over the ensembles (**Extended Data Figure 6e**). While iPAEs and average features were the most informative in their respective categories, enFoldX performance was highest when using the totality of our feature space, reiterating the importance of considering distributions of local and global structural metrics for this classification task.

### enFoldX outperforms sequence-based predictors and single structure-based methods

Most published TCR:pMHC prediction methods are based on natural language processing (NLP) models, like Large Language Models (LLMs), trained on the amino acid sequences of TCRs paired with cognate peptides^40–42^. A key limitation shared by all such models is the need for a large training corpus of functionally validated cognate pairs. By contrast, enFoldX, and other structure-based models, leverage protein structure prediction tools that are already pre-trained on structures from the Protein Data Bank (PDB)^43^ and, as such, require less TCR:pMHC specificity data for further training.

Because many published models cannot be retrained on new datasets, to systematically investigate the differences between sequence-only NLP models and structure-based ones, we developed BERTie (**BER**T for **T**CRs and **I**mmunogenic **E**pitopes) as a representative of the class of NLP models. Adapted from the widely used transformer architecture BERT (Bidirectional Encoder Representations from Transformers)^44^, BERTie is an LLM-based model trained exclusively on CDR3β and peptide sequences from single-chain TCRβ and peptide datasets (**Figure 4d**).

To optimize BERTie, we investigated several embedding regimes including single amino acid embedding (char) and *n*-gram embedding where input sequences are split into overlapping subsequences of length *n* (**Methods**). We tested each embedding regime, as well as a combined ensemble of char, 2-gram, and 3-gram, and found that the ensemble model achieved the best performance with an AUC of 0.87 for five-fold cross-validation (**Extended Data Figure 7a**). Evaluating BERTie on increasingly difficult classification tasks, we also found BERTie predicts for pairs of seen epitopes and TCRs with highest accuracy and performs substantially better than chance on pairs of unseen epitopes with unseen TCRs (AUROC=0.64, AUPRC=0.60) (**Extended Data Figure 7b**). We then benchmarked BERTie against several peer-reviewed best-in-class methods on the Immune Epitope Database (IEDB)^45^ and cross-reactivity^21^ datasets. BERTie performed as well as, or better, than other sequence-only models, some of which were trained on the CDRβ alone (ERGO-II^46^), others on paired ɑ/β chains (Tulip^42^, NetTCR2.2^47^) (**Figure 4e, f**). Notably BERTie’s performance characteristics mirror those of other NLP models, with relatively poor performance on the single amino acid substitution datasets and random performance on the full IMMREP25 dataset. This observation, along with BERTie’s strong performance compared with other NLP models, suggested that its generalization properties would reflect those of other models.

We then systematically evaluated different training-test set splits to study BERTie’s generalization capacity, finding that its classification performance is largely dependent on peptide and TCR sequence similarity to those included in training data (**Extended Figure 7c-f**). Based on this analysis, we conclude that sequence-only models can perform well for sequences close to the training data, but that structure-based models such as enFoldX are essential for generalization to TCRs and peptides that differ substantially from the pairs that they were trained on, or for distinguishing cognate and non-cognate peptides that differ by a single amino acid.

Our comparisons also reveal a clear advantage of enFoldX over sequence-based and other structure-based models. On our most challenging prediction benchmarks: the mutational scan validation sets and unseen data for IMMREP25, we scored methods for which code was publicly available (which excluded the top 8 structure-based methods submitted to IMMREP25)^18^. On the mutational scan benchmark, enFoldX outperformed all evaluated methods, including sequence-based models, the single structure-based models like TCRen, and a structure-based classifier which used the same feature set as enFoldX but only considered the highest AF3-ranked prediction (**Figure 4e**). The disparity between enFoldX and sequence-based predictive ability was further pronounced on IMMREP25 where sequence-based models largely show random performance (**Figure 4f**) with some variation on a per-epitope basis (**Extended Data Figure 8**). Exact comparisons with the top methods submitted to IMMREP25 were impossible, because the precise dataset used for scoring was not specified, so we instead scored enFoldX on the full, held-out test set and compared its AUROC at 0.1 false positive rate (AUC@0.1FPR) to that reported by IMMREP25 on the top methods. enFoldX outperformed all submitted IMMREP25 methods on median AUROC@0.1 FPR across 18 tested epitopes.

### Assessing the limitations of sequence- and structure-based TCR:pMHC predictors

Based on the mutational scan and IMMREP25 results, we aimed to quantify the limitations of sequence- and structure-based TCR:pMHC predictors, focusing on BERTie and enFoldX to represent each type of approach. For BERTie, we found that its accuracy depends on both epitope and TCR sequence similarity to training data with its most accurate predictions for epitopes at a Levenshtein distance of two amino acid substitutions from the training data (**Figure 5a**). While BERTie demonstrated excellent performance (AUC=0.93) for test epitopes within a distance up to two substitutions, its ability to generalize to novel epitopes significantly deteriorated as epitope sequences diverged further from those seen during training (AUC=0.58) (**Figure 5b**). Taken together, this suggests that over-reliance on training data sequence similarity inhibits BERTie’s ability to predict changes in TCR specificity due to subtle or divergent changes in the epitope. By contrast, enFoldX performance is largely independent of epitope sequence similarity (**Figure 5c**), explaining its improved scoring compared to BERTie for both the mutational scans with point-mutated peptides and IMMREP25 with unseen epitopes.

**Fig. 5.**
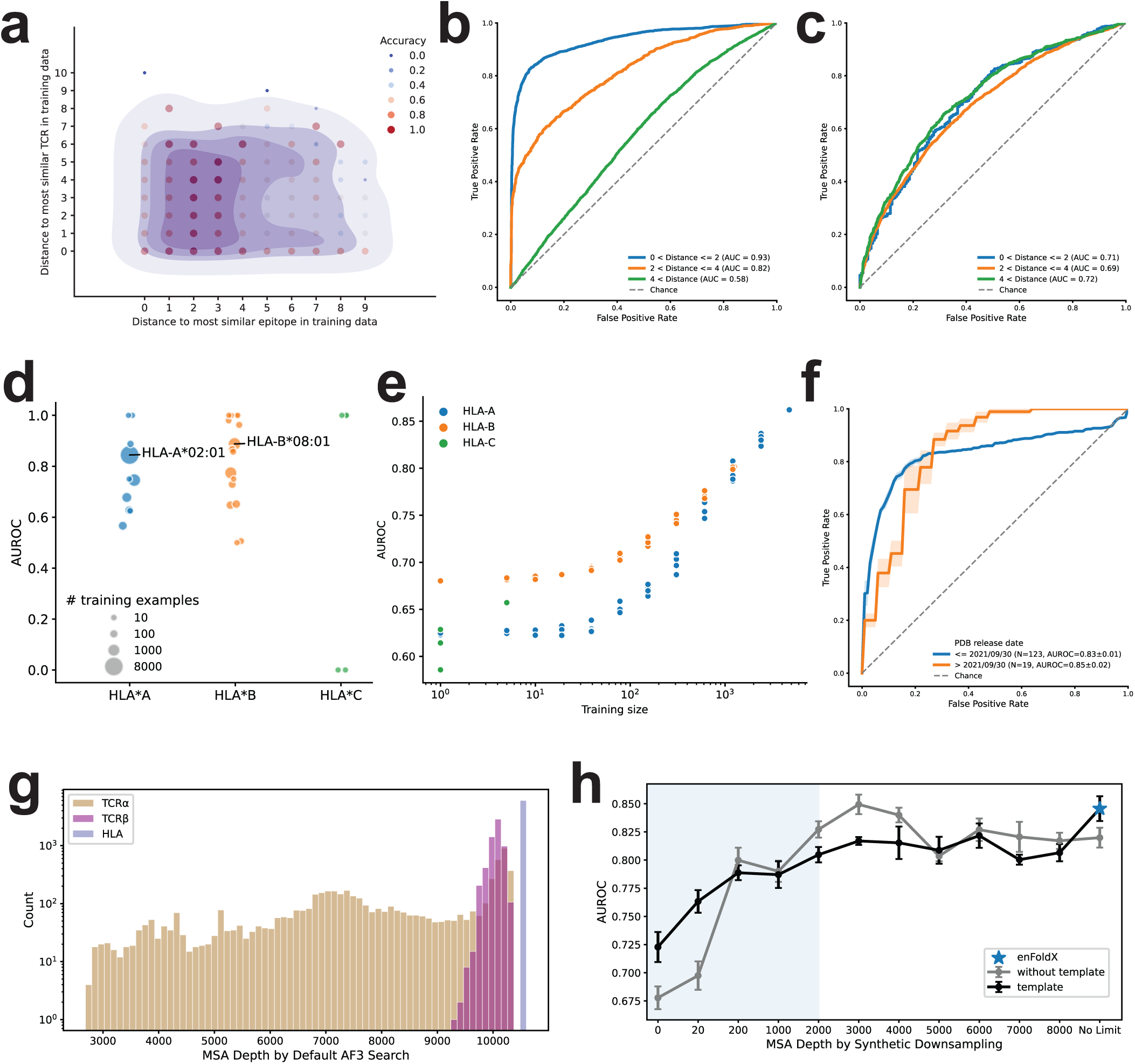
Assessing the limitations of sequence- and structure-based TCR:pMHC predictors. **a,** Accuracy of BERTie predictions as a function of similarity of TCR and epitope sequences to those in training data, measured by Levenshtein distance to the most similar sequence in training data. **b-c,** Performance on unseen epitopes based on distance to closest epitope in the training data for the sequence model BERTie (**b**) vs. enFoldX (**c**). The ROC curves for BERTie are based on training and test sets drawn from the union of TCRβ sequences from VDJdb+McPAS+COVID; those for enFoldX on training and test sets drawn from a subset of TCR:pMHC complexes from VDJdb 02/2025 for which paired TCR α and β sequences are available. The test set for BERTie are drawn from a 90/10 split stratified to include both unseen epitopes with seen TCRs and unseen epitopes with unseen TCRs. The test data for enFoldX is all epitopes in the training dataset evaluated with a ten-fold epitope-wise cross-validation. **d,** Performance of the enFoldX model on the VDJdb human decoy task by HLA allele and HLA-specific training set size. The model was evaluated using 10-fold TCR-wise cross-validation on the VDJdb 02/2025 human decoy dataset, and AUROC was computed separately for each HLA allele by pooling out-of-fold predictions across folds. Dot size is proportional to the number of cognate TCR:pMHC pairs available for each HLA allele. **e**, Performance of the enFoldX model on the VDJdb human decoy task by HLA group and training set size. To assess how much training data is required for generalization, the model was trained on the VDJdb 02/2025 human decoy dataset while systematically downsampling one HLA group at a time and evaluated on the corresponding HLA group in the independent VDJdb 09/2025 human decoy dataset. Each downsampling level was generated by random subsampling and repeated five times; each point represents one random subsample. **f**, enFoldX performance on TCR:pMHC complexes with published experimental structures, deposited in PDB before (blue) or after (orange) the AF3 training cutoff date. **g,** MSA depth distribution for individual TCR chains and the HLA alleles in the VDJdb 09/2025 data. **h**, Classification performance on the cross-reactivity dataset with randomly downsampled MSAs, including or excluding the structural templates in the folding inputs. Relevant MSA depth range based on **d** is shown in white background (above 2,000 sequences), with depths below that range in blue background. All models shown in panels b–f and h are enFoldX-Ridge models.

To better understand enFoldX’s limitations, we analyzed the impact of HLA representation in training data. Within TCR:pMHCs complexes with HLA-A and HLA-B alleles, there is no correlation between the amount of available data for a specific HLA allele and enFoldX performance, though AUC scores dramatically drop for certain HLA-C alleles (**Figure 5d**). We also observed a positive monotonic relationship between performance and training data size for all allele groups with AUC scores for HLA-B, the most polymorphic allele group, consistently outperforming the others as we systematically downsampled the training data for the corresponding HLA group (**Figure 5e**). This implies enFoldX’s stronger performance on HLA-B alleles is due to both the inherent diversity of the allele group and the scale of available data for training. By contrast, there is a clear deficiency in structural and TCR:pMHC specificity training data for HLA-C, limiting predictive ability.

We also evaluated enFoldX dependence on three factors crucial for high-quality AF3 predictions: class I TCR:pMHC structures incorporated in AF3 training; MSA depth, the number of homologous sequences returned by AF3 for a given query sequence; and structural templating^19,48^. To do so, we first matched class I TCR:pMHC structures in the PDB with VDJdb records, withholding these pairs from enFoldX training data. For classification of TCR:pMHCs with released PDB structures, we found no significant difference in enFoldX performance for TCR:pMHC structures seen or unseen in AF3 training, indicating independence from AF3 training data similarity (**Figure 5e**). We then analyzed AF3 outputs from its default MSA and template search for VDJdb data, determining the relevant range for per-chain MSA depth to be between 2,000 and 10,000 sequences with the most variability for TCRα chains (**Figure 5f**). Using our cross-reactivity dataset^21^, we re-ran the enFoldX ensemble generation step with artificially downsampled MSA, including or excluding structural templates selected by AF3. Performance by five-fold cross-validation remained consistent in the relevant MSA depth range, only decreasing steeply when the MSA depth was under 2,000 (**Figure 5g**). Notably, enFoldX still demonstrated predictive ability for TCR:pMHCs without any MSA or templates (AUC=0.68) when the structural quality of AF3 predictions was markedly poor (**Extended Data Figure 9**). Crucially, this indicates that even when AF3 structural predictions are clearly inaccurate, there are systematic differences between predicted cognate and non-cognate ensembles that enable customized enFoldX features to extract meaningful signal.

Investigating the varying accuracy in AF3 structural predictions, we revisited the predicted hallucinations presented in **Figure 1** for cognate and non-cognate TCR:pMHCs differing only by an epitope point mutation. We extracted subunit poses from the mutant non-cognate hallucination and re-aligned them to the predicted cognate wild-type structure (**Extended Data Figure 10**). While the overall re-aligned pose appears plausible, closer inspection of the non-cognate interface reveals strong steric clashes with virtually no distance between the mutant peptide side chain and CDR3β loop. As AF3 was trained on experimental structures and is therefore biased toward generating cognate models, we suggest that AF3 attempts to construct canonical TCR:pMHC geometry but, inferring unfavorable interfacial interactions that are explicitly penalized in AF3 scoring, predicts a non-physiological structure with the misdocked TCR. This implies that while the sampled conformational space of individual, unbound subchains can be limited by the AF3-imposed prior on co-evolutionary relationships, there may be more structural diversity sampled at the binding interfaces of multichain protein complexes.

## DISCUSSION

An ideal model for TCR specificity must perform well on two extremes: (1) novel pairings of unseen TCRs and pMHCs and (2) TCR:pMHCs that subtly differ by single point mutations. Published, state-of-the-art prediction methods are largely sequence-based with a strong dependence on sequence similarity to training data, limiting their generalizability^49^. By contrast, enFoldX demonstrates predictive ability on out-of-distribution TCR:pMHCs, as evidenced by our generalization from human to mouse data and our performance on IMMREP25 data, and TCR:pMHCs with single amino acid substitutions in the epitope, demonstrated by our mutational scan benchmark. We conclude structure-based ensemble approaches enable genuine transfer learning.

We attribute enFoldX performance to our consideration of the entire predicted structural ensemble and structural features relevant to TCR:pMHC binding. We have shown structural ensemble-based models improve classification compared to single structure- and sequence-based methods by capturing systematic differences in diffusion-based predictions for cognate and non-cognate TCR:pMHCs, including non-cognate pairs due to failed MHC presentation rather than failed TCR recognition. We have also demonstrated the utility of a diverse feature set spanning local and global measures with a particular emphasis on the TCR:pMHC interface with custom features for peptide-CDR3 interactions. Additionally, our approach is not limited to AF3. The general framework can be adapted as a downstream analysis pipeline for other structure prediction methods (Boltz^50,51^, RoseTTAFold^52^, OpenFold^53,54^) or protein language models like OmegaFold^55^ or ESMFold^56^. While we focus here on TCR:pMHC pairing with an AF3-based model, these lessons on ensembling structurally relevant features from protein structure prediction methods are broadly applicable to other *in silico* binding prediction challenges.

While we consider our enFoldX approach to be a significant advance in predicting TCR specificity, there remains substantial room for improvement, as predictions on unseen data remain challenging, motivating the need for more comprehensive, high-quality datasets of experimental TCR:pMHC structures, binding affinities, and activation. While structure-based methods showed gains in the IMMREP25 competition, those methods have been neither made available nor peer reviewed at the time of submission. Given the open and principled nature of our approach, and its extendibility beyond AF3, it provides an interpretable framework for identifying limitations in the underlying training data and for guiding future model improvements. For example, we can quantify deficiencies in available training data that limit better prediction, such as underrepresentation of specific HLA-C alleles, and limitations in the biophysical understanding of co-folding models that leverage co-evolutionary relationships to predict structures^57–59^.

We found ensembling to be of importance and a direction for future work, but we also acknowledge that it entails a significant increase in computational cost, which we quantify directly (**Extended Data Figure 2**). The power of ensembling lies in its ability to overcome a weakness shared by many diffusion-based models, like AF3. When applied to out-of-distribution data, these models often hallucinate, i.e., they confidently make random guesses, and ensembling reveals the disagreement among these guesses. Moreover, we suggest that our classifier, whose most important features were largely derived from customized points of biophysical contact, could capture a true free energy landscape. Ensembling therefore approximates the entropic favorability of the true minima over the many false minima along the way to protein folding^60^. Under the folding funnel hypothesis^61^, a rugged energy landscape demonstrates barrier ridges where misfolded intermediates fall in kinetic traps and fail to achieve their final conformation at the global minimum. Principled ensembling may therefore allow for a more robust sampling of this rugged space^60^.

Significant efforts have been made to collect TCR specificity data, though the most widely used databases such as VDJdb contain a very uneven distribution of TCRs across peptides, lack experimental examples of noncognate pairs, and may potentially contain a number of false positive records. While we benchmarked enFoldX and alternative methods on datasets with experimental positives and negatives, many others in the field have exclusively assessed performance with synthetic negative data, which can lead to inflated scores and an overestimation of generalizability^62^. We therefore argue that our field requires targeted, high-quality, publicly available datasets with experimental structures, binding affinity, reported negatives and T cell activation readouts, as well as greater consideration of biophysical properties governing TCR specificity in their generation. Doing so iteratively with enFoldX or other ensemble-based, biophysically grounded approaches will enable systematically more accurate, generalizable models, accelerating the development of precision immunotherapies. Advancing this paradigmatic protein binding prediction task will therefore both specifically advance public health goals, such as the development of vaccines and cell therapies against cancer, while also offering general lessons for AI in biology.

## ACKNOWLEDGEMENTS

We thank Alexander Solovyov, Cyrus Tam, Taykhoom Dalal, Zachary Azadian, and Lohit Valleru for helping with installing and running AlphaFold3. We thank Caleb Lareau, Hoyin Chu, Chrysothemis Brown, Gabriel Victora, Martina Milighetti, and David Hoyos for insightful discussions. We thank Nicole Rusk for editorial assistance. BG would like to thank John Hopfield for a helpful discussion. This work utilized resources from the High-Performance Computing (HPC) Group at Memorial Sloan Kettering Cancer Center. This work was in part supported by The Tow Foundation; The Marcus Foundation, and the Ben and Rose Cole Charitable PRIA Foundation (to V.P.B), the Center for Experimental Therapeutics at MSK (to V.P.B. and B.D.G.); an ASPIRE Award from the Mark Foundation (B.D.G.); Robert J. Kleberg, Jr. and Helen C. Kleberg Foundation (B.D.G. and Q.M.); and The Olayan Charitable Foundation. OL was in part funded by Memorial Sloan Kettering Cancer Center’s Center for Experimental Immuno-Oncology Scholars Program. JL is supported by NIAID Ruth L. Kirschstein National Research Service Award for Predoctoral Fellows 1F31AI200147-01.

## DATA AVAILABILITY STATEMENT

We provide easy access to the labeled mutational scan TCR-pMHC datasets used for validation in our github, at https://github.com/jonlevi/enFoldX. Other datasets used in the paper can be accessed directly from their original publications, as described in the methods below. The weights of our trained enFoldX ridge regression models are also included in our repository. The full set of predicted structure ensembles and extracted feature vectors can be made available upon request.

## CODE AVAILABILITY STATEMENT

We created a github repository with instructions and tutorials for running enFoldX at https://github.com/jonlevi/enFoldX. The repository includes instructions for how to use the tool, including tutorials both for a local installation of AF3 and for predictions using the AF3 Server. It also includes instructions on how to use enFoldX models that are pre-trained on VDJdb that can be used to predict for new TCR:pMHC sequence combinations.

## METHODS

### enFoldX

#### Structure Prediction Pipeline

Our pipeline involves three distinct steps which we run sequentially: structure look-up via template search and multiple sequence alignment (MSA), structure prediction via diffusion, and feature extraction from predicted structures and confidence metadata (**Figure 2**). We first collect all unique TCR α and β chains and MHC sequences in the dataset and run the first stage (“MSA”), where AF3 queries the input sequence against multiple sequence databases to find similar sequences and collects co-evolutionary information for structure prediction^63^. We omit beta-2 microglobulin (β2Μ) as it does not vary across complexes. We perform this step in a parallelized manner across all unique sequences. We then process the MSA output files and generate input files in JSON format for the second step: structure prediction. These JSON input files include TCRα, TCRβ, MHC, and peptides as four separate sequence chains, and MSA and template search results from the previous step for all chains except the peptide, which is generally too short to yield any meaningful MSA or template matches. We then run the protein folding step in parallel across all input files. For each input file, AF3 then produces multiple structures corresponding to the number of random seeds and the number of samples, which are set as hyperparameters in the AlphaFold input. The total number of predicted structures in the ensemble output will be *N* seeds times *K* samples per input. Unless otherwise specified, for the enFoldX models in this paper we chose *N*=10 and *K*=5 for ensembles of size 50 for both training set features and test set features across all the datasets.

#### Ensembling Features

Formally, the featurization of each TCR:pMHC maps an individual set of 4 input sequences (2 TCR chains, a peptide chain, and an MHC chain) to a vector of size 2*M*, where *M* is the number of features extracted from each structure, with one entry for the mean value of that feature across the ensemble and one entry for the standard deviation of that feature across the ensemble. This can also be thought of as modeling the distribution of features across the ensemble as a multivariate Gaussian distribution over the ensemble of an *M*-dimensional random vector.

#### Description of enFoldX Engineered Features

To build the representation described above for each of the *M* features, we first must compute and extract the values of that feature in each of the structures in the ensemble individually. For each of the 50 predicted structures per input, we leverage the “confidence” JSON files output by AF3, containing metrics that represent model confidence on the per structure level, per chain level, per residue level, per pair of chains, or per pair of residues.

enFoldX uses three types of base confidence metrics (pLDDT, pAE, pTM), and one type of structural metric (contact probabilities). As has been shown previously^24^, these metrics can be informative when extracted properly. Given that the TCR:pMHC complex has multiple interfaces, we included the average iPAEs between all pairwise groups of chains (α, β, MHC, and peptide), as well as some subchains (CDR3α, CDR3β) as features in our model, as described below. This iPAE is also sometimes referred to as interface PAE or interaction PAE and has mostly been leveraged in the context of predicting de novo proteins binding to protein targets^64,65^. Along with ipTM, these metrics have also been used as a general predictor of protein complex formation in place of traditional docking methods^66,67^. As will be shown below, we deviated from the standard approach in the literature and included other ways of collapsing these submatrices, including taking the minimum, maximum, or standard deviation of the entries. The contact probability between two residues indicates the predicted probability that their representative atoms are within 8Å of each other and is provided by AlphaFold3 as an output. We explored changing the threshold of 8Å to smaller distances using our own contact map extraction code, but the performance of enFoldX did not change significantly with alternative definitions of contact (data not shown).

The full feature set of size *M* is defined as follows:

For pLDDT and pAE, we compute:

i. The mean value over the global structure (= **2 features**)
ii. The mean, maximum, minimum, and standard deviation over values for each chain individually (4 chains times 4 aggregators times 2 metrics = **32 features**)

For pTM, we compute:

i. The mean value over the global structure (= **1 feature**)
ii. The mean over values for each chain individually (4 chains = **4 features**)

In addition to per-chain aggregations, we also compute inter-chain statistics of some of these metrics^2^, which we prefix with an “i”, such as ipAE and ipTM.

For ipAE, we compute:

i. The mean, maximum, minimum, and standard deviation over the PAE submatrix for each pair of chains (6 pairs times 4 aggregators = **24 features**)
ii. The mean, maximum, minimum, and standard deviation over the PAE submatrix for the TCR CDR3α/β regions paired with both the MHC and the peptide (2 CDR3s times 2 chains times 4 aggregators = **16 features**)

For ipTM, we compute:

i. The mean ipTM for each pair of chains (6 pairs = **6 features**)
ii. The mean ipTM value over the global structure (= **1 feature**)

For contact probabilities, we compute:

i. The mean and the maximum contact probabilities between each pair of chains (6 pairs times 2 aggregators = **12**)
ii. The mean and the maximum contact probabilities between TCR CDR3 regions (α and β) paired with both the MHC and the peptide (2 CDR3s times 2 chains times 2 aggregators = **8 features**)

These features make up the *M* dimensional vector, where ***M*=106**. The final feature set is thus **2*M*=212**, as each feature is extracted with a mean and a standard deviation across the ensemble as described above.

#### Pairwise RMSD Calculation

To quantify the diversity of a single ensemble of structures (as in **Extended Data Figure 1** and **Extended Data Figure 3**), we compute RMSD metrics (21 total) between all pairs of structures in the ensemble (i.e. for 50 structures, 502=1225 comparisons). For each structural pair, we first perform an overall alignment of the structures and compute RMSDs between the overall structures and each chain (overall RMSD + per-chain RMSD = 5 metrics). We then align the structures by chain and compute the RMSD on each chain (chain-by-chain RMSD = 16 measures). All superposition and RMSD calculations are performed using alpha carbon (Cα) atoms. These metrics can then be aggregated over the ensemble by mean, standard deviation, and min/max to generate an 84-feature vector describing the structural diversity of the ensemble. The script to compute pairwise RMSD is included in our GitHub repository. For visualization, the plots in this manuscript only feature RMSD over peptide Cα atoms.

#### Classification

From the curated feature set of size 2*M*, we trained classification models to distinguish between cognate and non-cognate complexes. For the within-data set classification, we performed ten-fold cross-validation by TCR and epitope with an L2-regularized logistic regression classification model (Ridge Regression)^31^, splitting folds by unique TCR or epitope sequences to mitigate data leakage between training and test data. The final ROC curves are the average across the folds. For validation datasets, we used VDJdb human data for training, with negative data as described in the text. For the cross-reactivity dataset, we also tested train/test splits using leave-one-out strategies for each position and each TCR. We also tested other architectures/penalty paradigms, and they achieved similar performance, with slight variability depending on the dataset, including XGBoost^38^ and Random Forest^39^ as implemented by scikit-learn^68^ (see **Extended Data Figure 6c**). We chose Ridge Regression for the bulk of the analyses since it can help alleviate multicollinear features (which is likely an issue given our feature construction process) and showed the best performance on the generalization tasks. For the IMMREP25 classification task, we observed an additional performance gain from combining XGBoost, Random Forest, and Ridge Regression into an ensemble model relative to Ridge Regression alone, and we therefore report results for each individual model and the model ensemble.

#### Feature Importance Analysis

To determine which features contributed the most to the classification, we conducted SHAP value analysis^32^ using the SHAP python package (https://github.com/shap/shap) for four scenarios: TCR-wise CV with decoy and permuted negatives and epitope-wise CV with decoy and permuted negatives. We calculated mean absolute example-level SHAP values for all features for each fold (in each of the four analyses), then computed the means across all folds for each analysis. Overall feature importances were very concordant between TCR-wise and epitope-wise CVs, but there were differences between feature importance in Decoy scenarios and in Permuted scenarios.

#### Evaluation of AF3 Training Data Structural Similarity

To annotate class I TCR:pMHC structures released in the PDB, we used TCR3d^69^, a TCR structural repertoire database. Annotations, including PDB IDs, chain sequences, and PDB release dates, were downloaded from TCR3b on 2026/02/01. Sequences from the collected PDB IDs and September VDJdb records were matched by epitope and CDR3α/β sequences and by HLA allele. Matched TCR:pMHCs were withheld from training data to retrain enFoldX on all September and February VDJdb data with scores equal to or greater than one, using both decoy and permuted negatives. Decoy negatives were constructed for complexes with available structures to evaluate whether enFoldX can discriminate real positives with available structures from decoy negatives (**see** **Figure 5f**).

#### MSA Depth Quantification and Subsampling Approach

We define MSA depth as the total number of homologous sequences returned by AF3 for a given query sequence. We computed MSA depth for all cognate TCR:pMHC pairs in our training set curated from VDJdb (6016 examples). By default, the enFoldX pipeline uses the full MSA depth and templates returned by AF3 for each chain. To evaluate the dependence of enFoldX performance on AF3 MSA and templating, we focused on TCR:pMHC pairs in the cross-reactivity dataset (1204 examples). After performing our default AF3 MSA and template search step and concatenating the results into AF3 folding inputs, we modified the inputs by setting a maximal MSA depth limit *m*. For chains with MSA depths greater than *m*, we randomly selected *m* sequences from the AF3 MSA search results to include in our folding input. To remove templates, we set the AF3 “templates” field for each chain to an empty list. We generated structures for the cross-reactivity dataset (5 seeds, 5 structures per seed = 25 structures per TCR:pMHC pair) with and without templates with twelve different maximal MSA depths, ranging from no limit to zero MSA depth, resulting in 24 datasets. We evaluated performance, indicated by AUROC score, with 5-fold cross-validation for (1) training and testing on each dataset and (2) training on the enFoldX dataset with default MSA/template search and testing on the other datasets (**see** **Figure 5h** **and Extended Data Figure 9b**).

#### GPU Cost and Runtime Analysis

enFoldX GPU runtime was tested on 6 different NVIDIA architectures. Profiling was done on a random sample of 1000 TCR-pMHCs from the mouse VDJdb dataset, and the runtime (seconds/per seed) is shown as a mean over 1000 TCR pMHC inputs, each with ten seeds. GPU rental cost estimates were obtained from the cloud GPU pricing aggregator GetDeploying Cloud GPU Index^70^, which compiles hourly pricing across multiple cloud providers. For simplicity, we used rental prices for AWS, a popular cloud provider. We extrapolated the per-seed estimates from the profiling above. For each architecture *a*, we computed the cost per seed as: Cost = Runtime * Hourly Rental Cost / 3600.

#### Structure Visualization

All structure visualizations were made using PyMol. In **Figure 1d**, **Figure 2**, and **Extended Data Figure 10**, structures are truncated to only show extracellular regions (peptide, MHC α chain, TCRα/β constant and variable domains) for visualization purposes. All other structures in this manuscript include the full-length sequences used in AF3 inputs. AF3 pLDDT coloring in **Extended Data Figure 10** was done using the following GitHub repository: https://github.com/cbalbin-bio/pymol-color-alphafold.git.

### DATASETS

#### Availability

All our curated labeled mutational scan datasets, along with the TCR, epitope, and MHC, are included in our github repository, at https://github.com/jonlevi/enFoldX. The other datasets used in the paper can be accessed directly from their original publications, as described in the methods below. The actual predicted ensembles of structures and the distributions of features can be made available upon request.

#### Stitching TCR Sequences

For input to the pipeline, we use full length TCR sequences. To do this, we used Stitchr^71^ to stitch the TCR sequences from the V and J gene and CDR3 information and filtered out any rows for which Stitchr failed or inserted a stop codon. A tutorial for how we use this tool and examples for running it are included in our github repository.

#### MHC Sequences

For mouse data, the amino acid sequences for the MHC molecules were downloaded from Uniprot^72^, with the data for H-2-Db from https://rest.uniprot.org/uniprotkb/P01899.fasta and for H-2-Kb from https://rest.uniprot.org/uniprotkb/P01901.fasta. For all human datasets, HLA sequences were obtained from the IPD-IMGT/HLA^73^ at https://www.ebi.ac.uk/ipd/imgt/hla/alleles/. For convenient access, all MHC sequences used in any dataset in our manuscript are included as fasta files in our github repository.

#### Quadratic Entropy

We calculate quadratic entropy using Rao’s formula^74^:

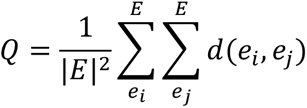

where *E* = {*e_i_*} is the set of epitopes in the dataset, and the distance metric *d* is calculated using the Levenshtein distance of amino acid sequences. This formula can be thought of as computing the expected Levenshtein distance between two epitopes drawn at random from the dataset.

### VDJdb DATASETS

#### Mouse VDJdb Data

VDJdb^28^ version 2025/02/21 was downloaded from vdjdb.cdr3.net. For the mouse data set, we filtered the data to ‘species’==’MusMusculus’ and only kept rows with non-null TCRs for both chains, non-null MHC, and a non-null peptide. We further filtered and kept only data for the MHC-I alleles H-2-Kb and H-2-Db. We used Stitchr^71^ to stitch the TCR sequences from the V and J gene and CDR3 information and filtered out any rows for which Stitchr failed or inserted a stop codon. This yielded 1896 positive TCR:pMHC combinations, 718 of which were for H-2-Kb (38%) and 1178 of which were for H-2-Db (62%), corresponding to 24 unique peptides.

To create negative pairs, we tried two alternative approaches. (1) For the decoy task, we matched each of the 1896 rows in the data with a decoy row, where the peptide was swapped out with a random decoy peptide from IEDB^45^ that was predicted by netMHC-4.0^29^ to be a strong binder to the MHC allele for that row. Peptides were downloaded from IEDB (mar10_2025.csv version) and filtered to include peptide sequences between 8 and 12 amino acids in length with “Object type”=”linear peptide”. Peptides that were already included in our VDJdb dataset were filtered out.

We defined binders as having Rank < 0.5 as is standard. (2) For the permutation task, we matched each of the 1896 rows in the data with a permuted negative where the peptide was swapped out with a different peptide from the dataset, where the other row had a different TCRα and TCRβ but the same MHC. In principle, this means that the peptide was effectively presented on that MHC to a different TCR but has no prior relationship to the TCR it was matched with and thus can be assumed to be non-binding.

#### Human VDJdb Data (February)

As described above, version 2025/02/21 (which we refer to as 02/2025 on plots) was downloaded from vdjdb.cdr3.net and filtered to include only the rows with ‘species’=’ HomoSapiens’ and entries that contain TCR α and β chains, a valid human MHC class I allele, and a non-null peptide. As with the mouse dataset, we used Stitchr^71^ to generate full-length TCR sequences from the V gene, J gene, and CDR3 information for α and β chains, and filtered out any rows for which Stitchr failed or inserted a stop codon. We further filtered the dataset to records where VDJdb score was 1 or above and agreed for both TCR α and β chains. This yielded 6016 positive TCR:pMHC combinations. We then followed the same strategy as described above to create the negative (non-binder) examples, either via the decoy approach or the permutation approach. For the decoy task, peptides were again downloaded from IEDB (mar10_2025.csv version) and filtered to include peptide sequences between 8 and 12 amino acids long with “Object type”=”linear peptide”; peptides that were already included in our VDJdb human dataset were filtered out. MHC presentation for each of the MHC alleles in the VDJdb Human dataset was predicted using NetMHCPan4.1^30^, and peptides predicted as “Strong Binders” were substituted for each positive TCR:pMHC example to create the decoy negative TCR:pMHC examples. For the Permutation task, the negative examples were created as described above.

#### Human VDJdb September

We downloaded version 2025/09/25 (which we refer to as 09/2025) from vdjdb.cdr3.net and filtered to include only the rows with ‘species’=’ HomoSapiens’ and entries that contain TCR α and β chains, a valid human MHC class I allele, and a non-null peptide, and VDJdb score of 1 or above. We processed the TCR sequences with Stitchr^71^ as described above and created decoy and permuted negatives following the same strategy as described above. For analysis, we considered any data deposited in VDJdb September that was not part of our VDJdb February dataset as new, yielding 1,681 new records.

#### TCRvdb

We obtained the copy of TCRvdb^33^ dated 01/05/2025 from the authors following the request protocol on the github repository, at https://github.com/schumacherlab/TCRvdb. This database consists of experimental re-validation data for TCR:pMHC complexes with 2 peptides: YLQPRTFLL and GLCTLVAML. We considered records with the adjusted p-value <0.05 as re-validated and therefore cognate, and records not meeting this threshold as non-cognate. 427 complexes from TCRvdb matched our filtered VDJdb 02/2025 dataset, with 259 complexes revalidated as cognate and 168 as non-cognate. For the analysis in Figure 4, we excluded all matching complexes from our training data, retrained enFoldX on VDJDB 02/2025 records excluding those found in TCRvdb, and predicted cognate/non-cognate status for these 427 complexes, achieving the AUC of 0.76. This dataset is not included publicly in our repository and should be requested directly from the authors as directed in their github linked above.

### OTHER DATASETS

#### IMMREP25 Test Data

The IMMREP25 competition^18,37^ contained a novel dataset from Adaptive Biotechnologies of 10,000 records with 1,000 TCRs paired with 20 virally-derived peptides presented by one of two MHC molecules (HLA-A*02:01 and HLA-B*40:01). All peptides are four or more Levenshtein distance from any published peptide and share no substring longer than five consecutive amino acids with any published peptide. The data was downloaded as linked in the pre-print from figshare^75^. For the IMMREP25 competition and results reported in the preprint, the publicly available test set was then randomly downsampled and subsequently filtered by the authors to remove leakage-prone cross-split records. The resulting IMMREP25 evaluation set (7344 records) was not released and could not be reconstructed. Therefore, we report results for the full published 9,000 records pertaining to 18 pMHCs included in the private IMMREP25 leaderboard without downsampling or filtering.

#### 1E6 TCR Data

Epitope data for this TCR was sourced from Wooldridge et. al.^35^, from Supplementary Table S3 (N=185 peptides). TCR sequences were downloaded from the PDB entry 5C0A. As the paper shows, this TCR has promiscuous binding to nearly every peptide with the central GP{D/F} motif. Thus, synthetic negative data was created by creating one negative decoy peptide for each positive peptide, where each amino acid in the central GP{D/F} motif was swapped to a random amino acid.

#### MEL TCRs Data

Data was sourced from Dolton et. al.^36^ (Figure 2, Figure 3, and Table S1). Data is shown for all three positive epitopes and six of the negative peptides that were used as irrelevant tetramers or shown to be false positive cognates from the “CANTiGEN” screen in Figure 2. All epitopes are presented on and modeled with HLA-A*02:01.

### PEPTIDE MUTATIONAL SCAN DATASETS

We collected a variety of mutational scan datasets to validate our model. These experimental binding/activation datasets start from a known TCR:pMHC pair and mutate the peptide at every position to every other possible amino acid, and then experimentally assay whether the TCR can still bind the mutated peptide. The datasets we included spanned a variety of experimental methods, including binding, activation, and killing assays. Most assays also do not use varying concentration sweeps but rather choose a single peptide concentration throughout, and each assay may vary in concentration the used. This can impact the resulting readout of the experiment and can mask the importance of MHC presentation on a positive experimental result. We only included datasets that did complete mutational scans (with the sole exception of TIL1383l which excluded anchor positions 2 and 9). We included datasets that had either raw data available, or had data with a clear normalization strategy, and that had a reasonable, well-defined threshold to define positives vs. negatives. Each one is described below in detail.

#### Cross-Reactivity Data

The T-cell “cross-reactivity” dataset is publicly available as part of our previous paper^21^. The half maximal effective concentration (EC50) fitted to the measured avidity curve for each of the 1204 TCR:pMHC combinations was binarized so that TCR:pMHC complexes with EC50 greater than 0.1 μg/mL were considered as non-reactive (non-binders), using the same threshold as in the original publication. This resulted in 342 binder TCR:pMHCs and 862 non-binders, across 7 TCRs.

#### NY-ESO Data

This data includes 3 TCRs with a mutational scan of the NY-ESO peptide. While the dataset was originally from Coles et al.^76^ (see Figure 4D-F there), we downloaded the data from the BATCAVE database^77^ at https://github.com/meyer-lab-cshl/BATMAN-paper/tree/main/results_batman/tcr_epitope_datasets/mutational_scan_datasets/TCR_data_input_by_publication/MHCI/NYE-Sx_TCRs. Because the original publication ran the mutagenesis assay with a single concentration (10 μM peptide), and did not do a concentration sweep, it was not possible to get a functional EC50 for every peptide. We thus took the average value of the raw IFNγ ELISpot across the 3 replicates and calculated the relative value for each mutant compared to the WT at that concentration. We set a stringent threshold of relative value >= 1 for strong binder cognates, corresponding to the middle of the color scale in the figure in the original publication.

#### PTEAM Data

Data was sourced from the PTEAM paper^27^, which is a mutational scan of the CMV epitope NLVPMVATV (Data S4 in the PTEAM paper). For the PTEAM data, activation scores were normalized based on controls as described in the original paper. We used the binding threshold recommended by the authors as laid out in their publication. Notably, this threshold is different from what is used for the same source dataset in the BATCAVE paper^77^, which used an alternative one-size-fits-all normalization strategy on all of the datasets they collected.

There is an additional mutational scan dataset in the PTEAM paper for a neo-epitope (VPSVWRSSL), but this scan excludes mutations across a variety of positions that lead to a loss of predicted MHC binding using in-silico predictions and only kept ∼75% of possible mutants. We chose not to include this dataset in our analysis, as it was the only one that used peptide presentation predictions to filter the scan, and we could not determine how this filtering would affect our ability to validate our model on this task.

#### Minigene Data

This 2 TCR mutational scan dataset (with the LLFGYPVYV peptide) was originally produced using a novel minigene depletion method in the PresentER paper^78^. We downloaded the data from the BATCAVE database^29^ at https://github.com/meyer-lab-cshl/BATMAN-paper/tree/main/results_batman/tcr_epitope_datasets/mutational_scan_datasets/TCR_data_input_by_publication/MHCI/TCRs_A6_B7. As before, we set a stringent threshold of relative depletion <= 0.5 for strong binder cognates, corresponding to the middle of the color scale in the figure in the original publication.

#### T1D Data

The mutational scan of the autoreactive 1E6 CD8+ T cell clone with the ALWGPDPAAA peptide was sourced from Figure 2 in Bulek et. al.^79^. This is the same TCR we analyzed in the 1E6 section.

#### MAGE Data

This dataset was sourced from a paper that developed the TCR-MAP method to study the targeting of the cancer associated MAGE-A3 epitope^80^. This method uses a synthetic circuit to activate sortase-mediated tagging of engineered cells expressing pMHCs. Figure 6B of that paper shows peptide fold enrichment data using TCR-MAP and was downloaded from the “Source Data” tab. Notably, this engineered TCR has been known to cross-react with an unrelated epitope from the Titin protein presented on cardiac tissue^81,82^. This dataset is also included in the BATCAVE repository. As before, we set the threshold more stringently than in BATCAVE, opting to use the approximate halfway mark from the original publication as a stringent threshold for strong binders.

#### SARS-CoV-2 Data

This dataset (which is not part of the BATCAVE dataset) was sourced from Figure 3 and accompanying source data from Malone et. al.^83^, containing mutational scans of the SARS-CoV-2 epitope YLQPRTFLL. The binding threshold was set to 1 as is used by the authors of that manuscript (see Figure 3d).

#### TIL1383l Data

This dataset was originally generated by Singh et al.^84^ to investigate T cell activation and TCR:pMHC binding with wild-type epitope YMDGTMSQV presented by HLA-A*02:01 and recognized by class-mismatched TCR TIL1383I originally cloned from a CD4+ T cell. A notable feature of TIL1383I is an exceptionally long CDR3β loop. We downloaded the raw data from the BATCAVE database https://paperpile.com/c/CNWfaF/XRFV at https://github.com/meyer-lab-cshl/BATMAN-paper/tree/main/results_batman/tcr_epitope_datasets/mutational_scan_datasets/TCR_data_input_by_publication/MHCI/TIL1383I. To binarize binding ability (originally measured using the UV-mediated peptide exchange assay data presented in Figure 5b from Singh et al.), we used K_D_ values calculated in BATCAVE and then set the binding threshold for TCR:pMHC pairs at K_D_ ≤ 200 μM.

### LARGE-LANGUAGE MODEL (BERTie)

#### Architecture

BERTie model consists of a BERT encoder and a Multilayer Perceptron (MLP) layer. For each CDR3β-epitope pair we prepend a [CLS] token and concatenate the two sequences together adding a [SEP] token between them and at the end. The sequences are padded on the right to maximum length within each batch. The BERT encoder takes in the tokenized sequence <t0, t1, t2, … tT>, where t0, t1, … represent amino acids in CDR3β or epitope or the special characters, and T denotes the index of the last token in the combined sequence. The model then generates the hidden representation from the last layer for each token of the input sequence: fE: ti → hi, ∀i ∈ {0, 1, 2 … T}. We then pass the hidden state of the [CLS] token hCLS to the Multi-Layer Perceptron (MLP) classifier layer which makes a prediction fD: hCLS → [0,1], where 1 indicates that the pair is predicted to be valid (cognate) or 0 – invalid (non-cognate). A weighted binary cross-entropy loss evaluates how close the predictions are to the truth and is back propagated through the model to update weight parameters in all previous layers in the direction that would decrease this loss. We developed BERTie versions where tokens are single amino acids and combinations of 2 or 3 amino acids. Where we refer to the BERTie ensemble, predictions of BERTie models based on each encoding approach are ensembled (averaged) to arrive at the ensemble prediction.

#### Datasets for BERTie

BERTie is trained and evaluated on three publicly available datasets: Adaptive Biotechnologies’ ImmuneCODE database containing putative SARS-CoV-2-specific TCR sequences and their corresponding epitopes (MIRA)^85^; VDJdb^28^ (described above); and McPAS-TCR^86^, a manually curated database of TCR sequences associated with various pathologies and antigens.

ImmuneCODE/MIRA contains 143,938 unique CDR3β sequences from TCRs of over 1,400 subjects exposed to or infected with SARS-CoV-2 virus. In a series of multiplexed experiments based on MIRA assay technology, these TCRs were co-cultured with various segments of SARS-CoV-2 viral peptidome and were found to pair with 363 unique SARS-CoV-2 epitopes. For BERTie we used VDJdb version 2022-03-30, a compilation of 51,257 sequences of TCRβ CDR3s from different studies and 227 epitopes known to be recognized by these TCRs (there are more records than used for enFoldX training, since BERTie requires only CDR3β sequences and not paired α and β TCR chains). McPAS contains over 20,000 TCR sequences matching over 300 unique epitopes with minimal overlap with VDJdb. The three datasets were de-duplicated, and only CDR3s of TCRβ chains for human MHC class I examples were used as input. The described datasets only contain positive TCR-epitope pairs. We generated negative examples by creating permutations of TCR and epitope sequences to create pairs not seen in the dataset.

BERTie was trained on 80% of the combined dataset and tested on the remaining 20%; however, where indicated in the main text, another version of BERTie was then retrained on all of the human MHC class I data.

BERTie was trained on single chain TCRβ and peptide datasets and used only the CDR3 portion of the TCRβ chains. We carefully created the test dataset so that it included equal proportions of 4 progressively more challenging test case scenarios: a new pairing of a TCR and an epitope both seen during training; a new (unseen) epitope paired with a TCR seen during training; an unseen TCR paired with a seen epitope; and a new pairing of an unseen TCR and an unseen epitope. We investigated several embedding regimes, including single amino acid embedding (char), 2-gram embedding where the input sequences are split into overlapping subsequences of length 2; 3-grams; and 4-grams. We then created an ensemble of 3-gram, char, and 2-gram models which achieved the best performance of 0.87 AUROC (**Extended Data Figure 7**).

BERTie was benchmarked against other sequence methods on the IEDB dataset^45^. IEDB TCR specificity dataset was downloaded on April 24, 2023, filtered to human TCRs with valid CDR3β, V gene, J gene, and epitope annotations, and deduplicated against VDJdb, McPAS, and MIRA. The final benchmark consisted of 2,186 unique positive TCR–epitope pairs and 2,186 permuted negative pairs.

### MODELS USED FOR BENCHMARKING

#### TULIP

TULIP^42^ is an unsupervised transformer-based model that learns TCR–epitope interaction patterns directly from positive binding pairs without requiring synthetic negative examples. By avoiding negative-sampling biases, it is designed to generalize more effectively to unseen epitopes and demonstrated state-of-the-art performance on multiple TCR specificity benchmarks when it was released (pre-IMMREP25). Notably, it was not tested on cross-reactivity datasets. TULIP was run using the pre-trained model and code example deposited at https://github.com/barthelemymp/TULIP-TCR.

#### NetTCR 2.2

NetTCR 2.2^47^ is a deep-learning model that predicts TCR–peptide specificity by combining peptide, HLA, and paired TCRαβ sequence information with large-scale public TCR dataset. We installed the code from their public repository at https://github.com/mnielLab/NetTCR-2.2 and ran their pre-trained pan-peptide model.

#### ERGO II

ERGO-II^46^ (referred to as ERGO on plots) comprises multiple neural-network architectures that encode TCR and peptide sequences using either LSTM or autoencoder representations and classify binding using a downstream multilayer perceptron. We ran the web version of ERGO II with all 4 available combinations: Autoencoder (AE) or LSTM based model trained on VDJdb or McPas datasets. We are reporting the best performing model, which was the AE model trained on VDJdb.

#### TCRen

TCRen was run using the R version of the TCRen code deposited in https://github.com/antigenomics/tcren as described in the TCRen paper^11^. For the mutational scan data, TCR:pMHC complexes with the WT peptide were modeled using homology modeling from TCRpMHCmodels-1.0^87^, as directed by the TCRen paper. Models were generated using the web server at https://services.healthtech.dtu.dk/services/TCRpMHCmodels-1.0/. All mutant peptides for the scan were then scored as “candidate epitopes” using run_TCRen.R with the default TCRen_potential.csv. The energy scores were then transformed as -1*Energy for classification. Note that for the IMMREP25 competition, the TCRen entry was updated to use a fine-tuned version of AF3 as outlined in the methods of the pre-print^18^, and we report their score for that dataset based on the released competition results. However, as this structure prediction method is not publicly available, we used the homology-based method recommended in the original TCRen publication for our direct comparisons on the mutational scan datasets.

## EXTENDED DATA FIGURES

**Extended Data Fig. 1.**
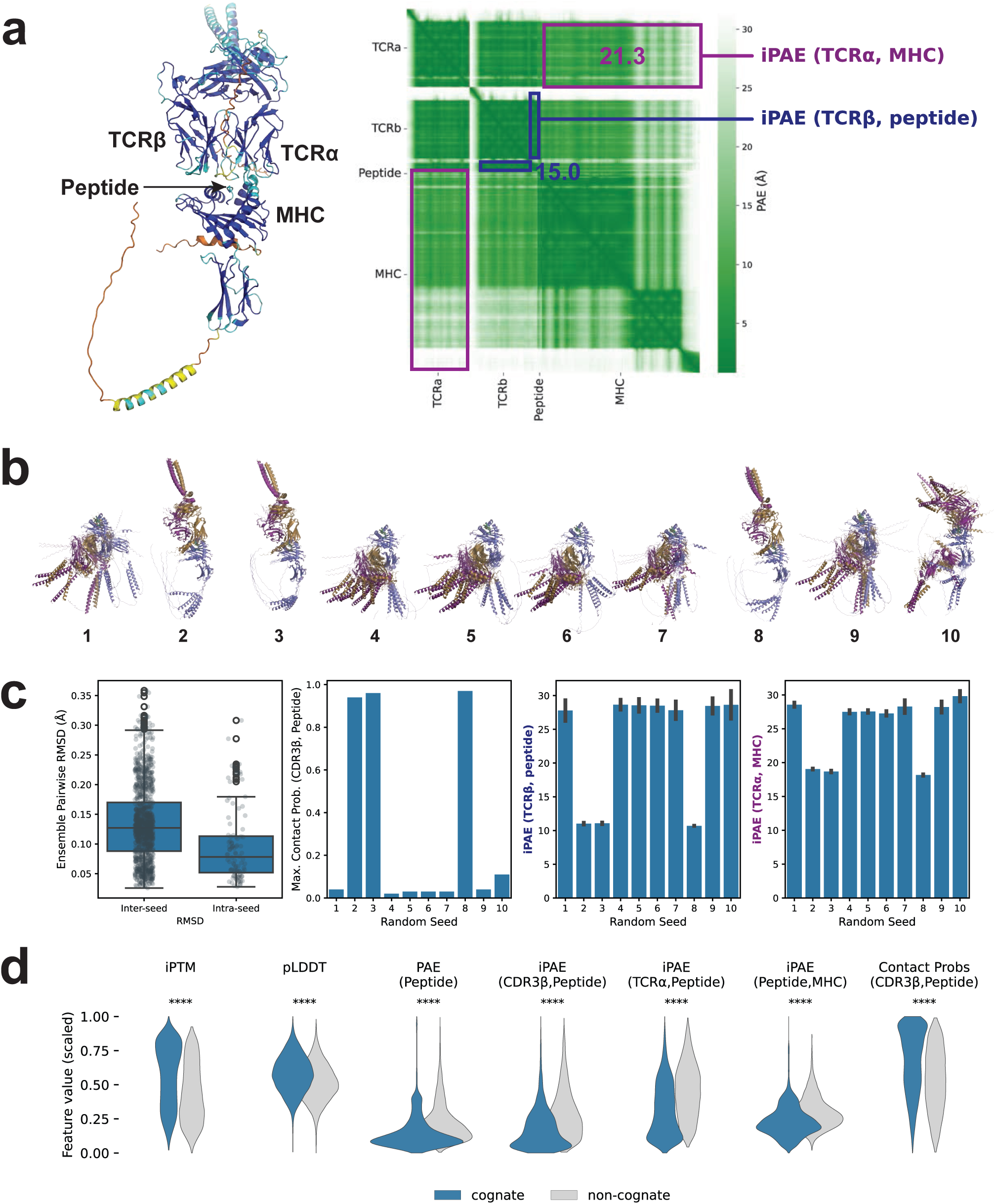
Ensembles of structures and confidence distributions for TCR:pMHC complexes. **a,** Example predicted AF3 structure for an experimentally validated binder, colored by pLDDT, along with the corresponding predicted alignment error (PAE) matrix. Annotations of the PAE matrix illustrate interfacial iPAEs (blue: peptide and TCRβ; purple: MHC and TCRɑ). Means of the PAE submatrices are annotated in corresponding colors. Higher iPAEs suggest lack of confidence in the predicted structure and may potentially reveal unfavorable binding. **b,** Predicted ensemble for TCR5-2 in complex with “non-cognate” M5K peptide from Figure 1d. Each number corresponds to five aligned structures sampled for an individual random seed (1-10). Visual inspection shows that structures vary more across seeds than samples. **c,** Diversity of the ensemble comes more from incorporating additional random seeds (inter-seed) rather than additional samples within the same seed (intra-seed) as quantified by pairwise RMSD, CDR3β-peptide contact probabilities, and interfacial iPAE distributions. **d,** Representative ensemble features related to confidence (ipTM, PAE, iPAE) and interchain contact probabilities are significantly different for cognate (positive) versus synthetic non-cognate (negative) TCR:pMHC complexes. Feature distributions are plotted for the VDJdb 02/2025 dataset (cognate, in blue) and for the corresponding synthetically created decoy negatives, as described in Methods (non-cognate, in gray). **** indicates p < 0.0001 by a two-sided t-test.

**Extended Data Fig. 2.**
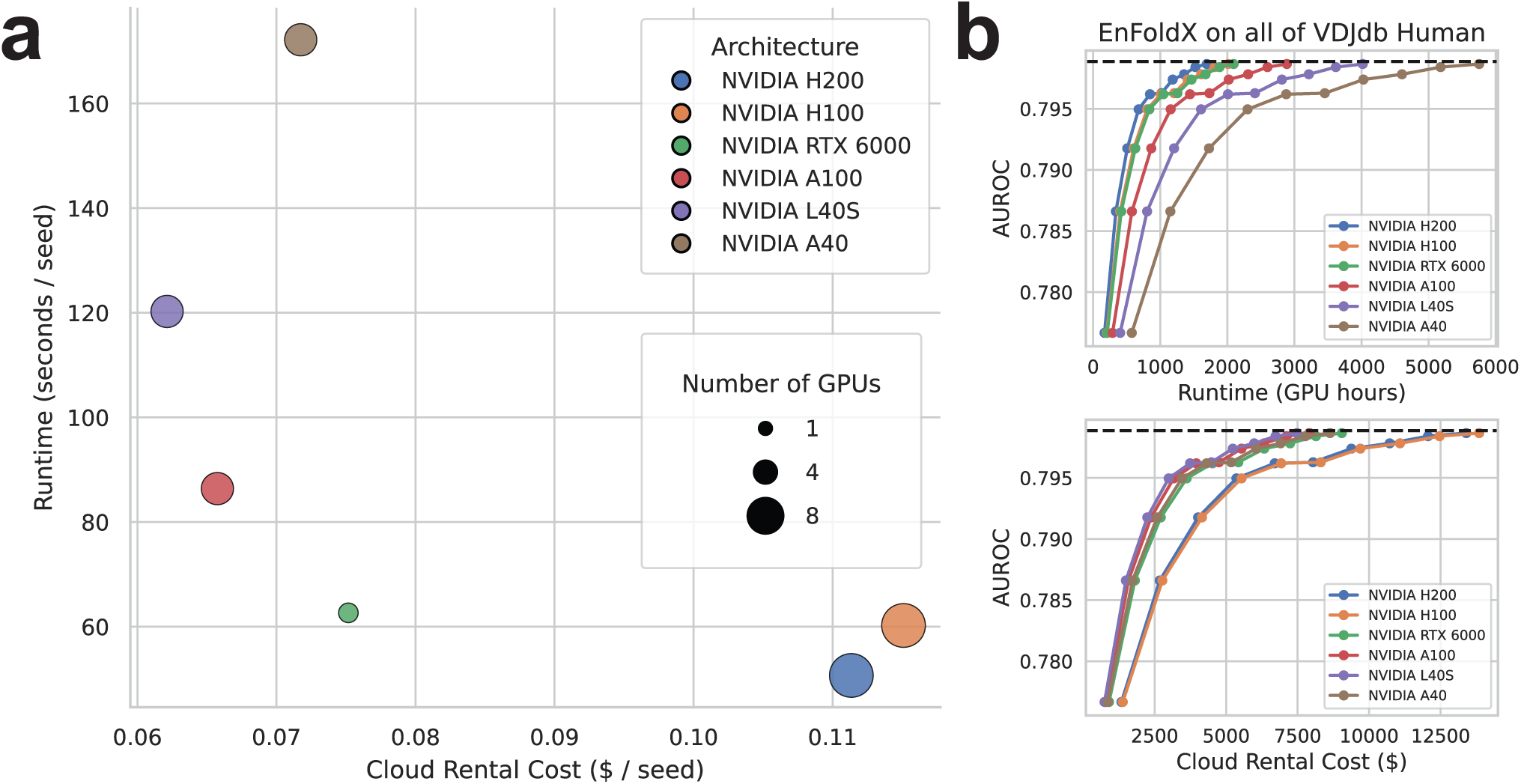
Profiling runtime and cost for enFoldX pipeline. **a,** Cost-per-seed vs. runtime-per-seed to run enFoldX for a single TCR:pMHC on six different GPU architectures (**Methods**). **b**, Tradeoff between runtime (top) or cost (bottom) and performance for running enFoldX with additional structures on VDJdb human data. Each dot shows the addition of another random seed in the AF3 ensemble, with AUROC performance numbers as in Fig. 1b.

**Extended Data Fig. 3.**
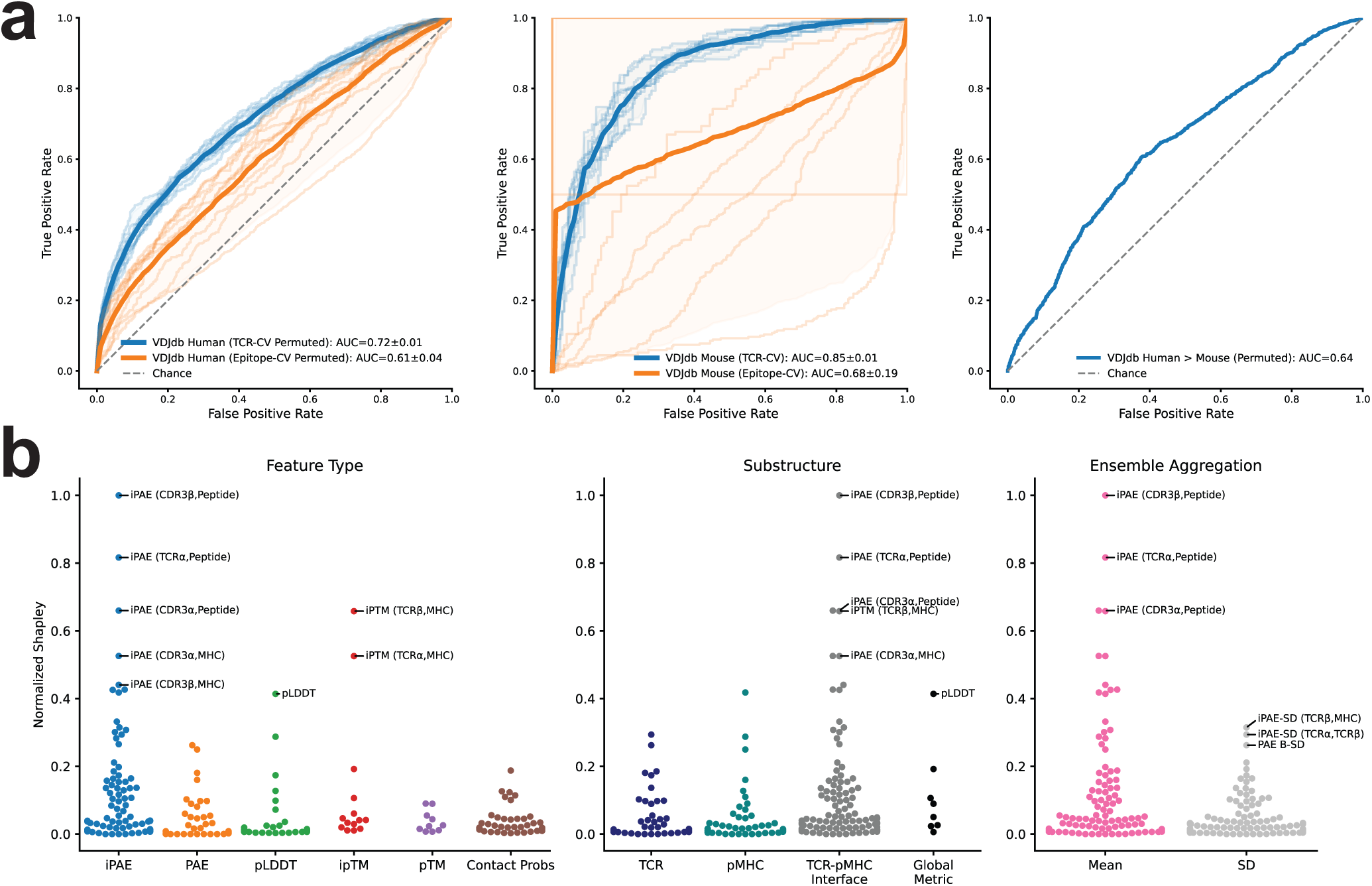
Predicting TCR:pMHC specificity for data in VDJdb with permuted negatives. **a,** ROC curves for classification of positive vs. permuted negative TCR:pMHCs in human (**left**) and mouse (**center**) VDJdb 02/2025 datasets, and for cross-species generalization (**right**). For individual human and mouse datasets, AUROC values are shown as mean and confidence intervals over folds using ten-fold cross-validation, splitting folds based on unique TCR (blue) or epitope (orange) sequences to mitigate data leakage between training and test data. For cross-species generalization, performance is shown for classification of TCR:pMHCs in mouse VDJdb using an enFoldX model trained exclusively on human VDJdb data**. b,** Normalized feature importances by SHAP (**Methods**) for the human VDJdb classification task with permuted negatives shown on the left in **a**. Features are stratified by feature type (**left**), substructure (**center)**, or by ensemble summary statistic (**right**). All results are for enFoldX-Ridge models.

**Extended Data Fig. 4.**
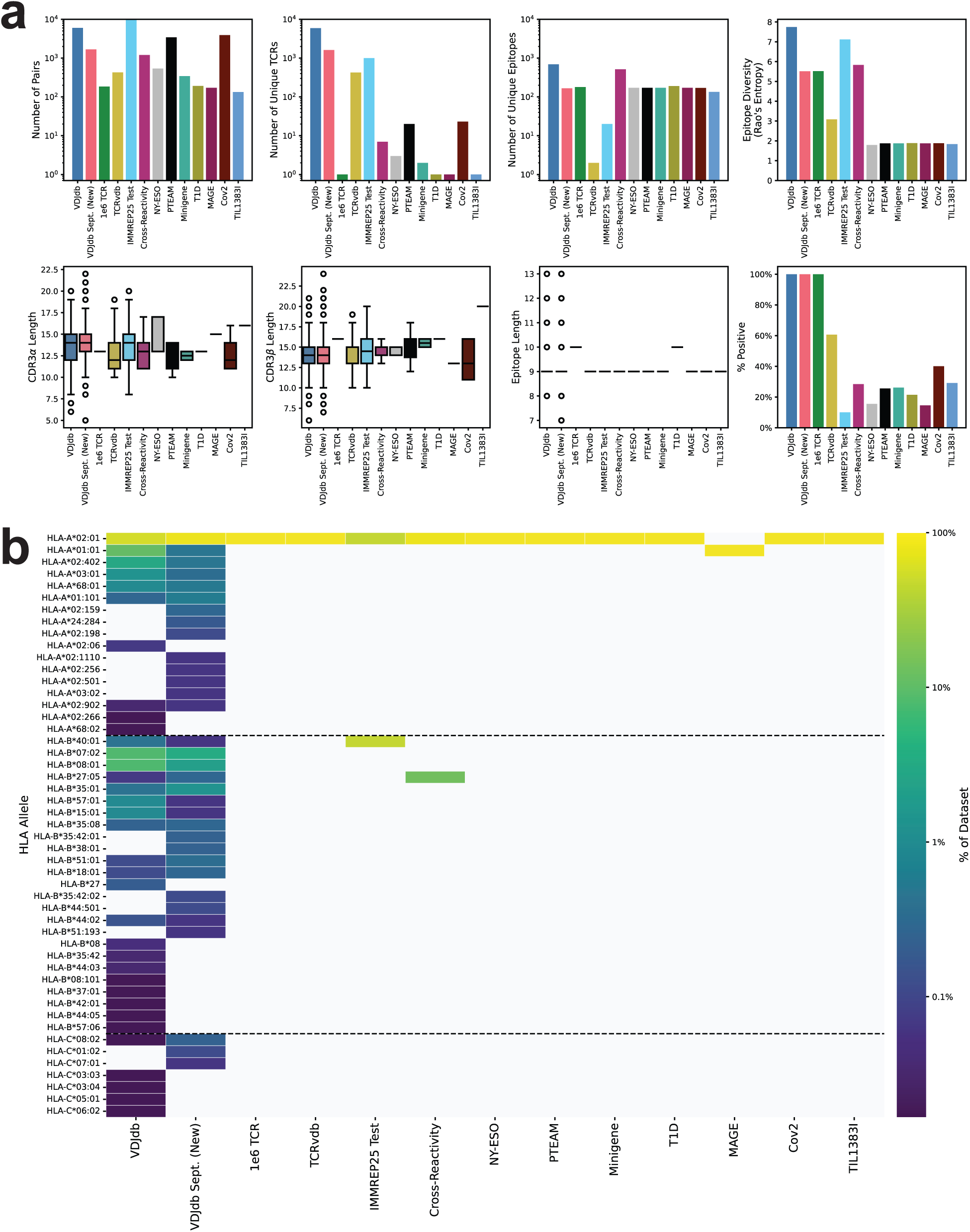
Summary statistics of validation datasets. **a,** Summary statistics for all experimental data used as validation in this paper. Data is only shown for experimentally validated data; synthetic negatives are not shown. For details on the datasets, see **Methods**. **b,** HLA representation across validation datasets; color indicates the percentage of TCR:pMHCs with the specified HLA allele.

**Extended Data Fig. 5.**
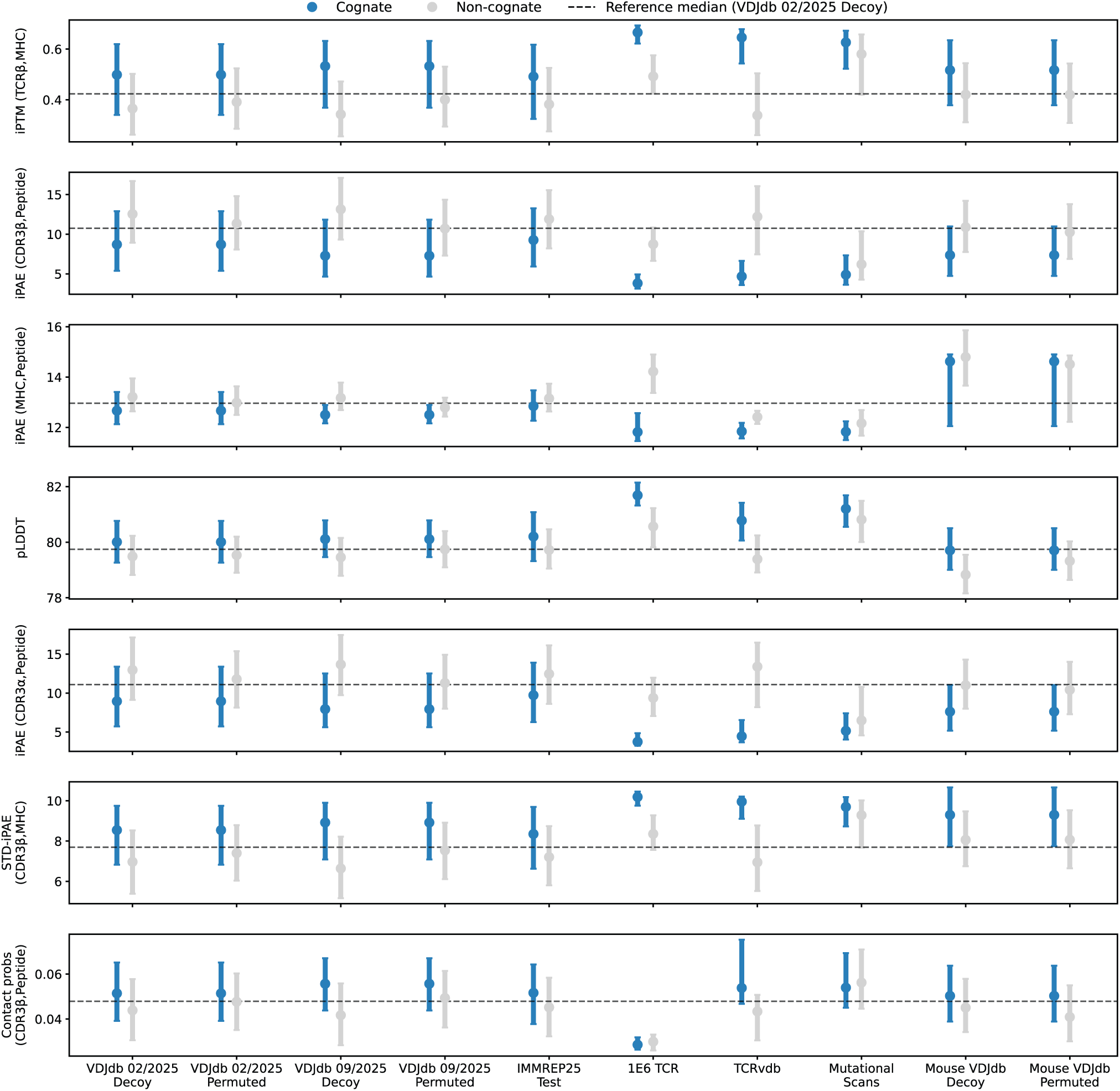
Distributions of example enFoldX features. Distributions of example enFoldX features for cognate (blue) and non-cognate (gray) TCR:pMHCs across the datasets used in the paper.

**Extended Data Fig. 6.**
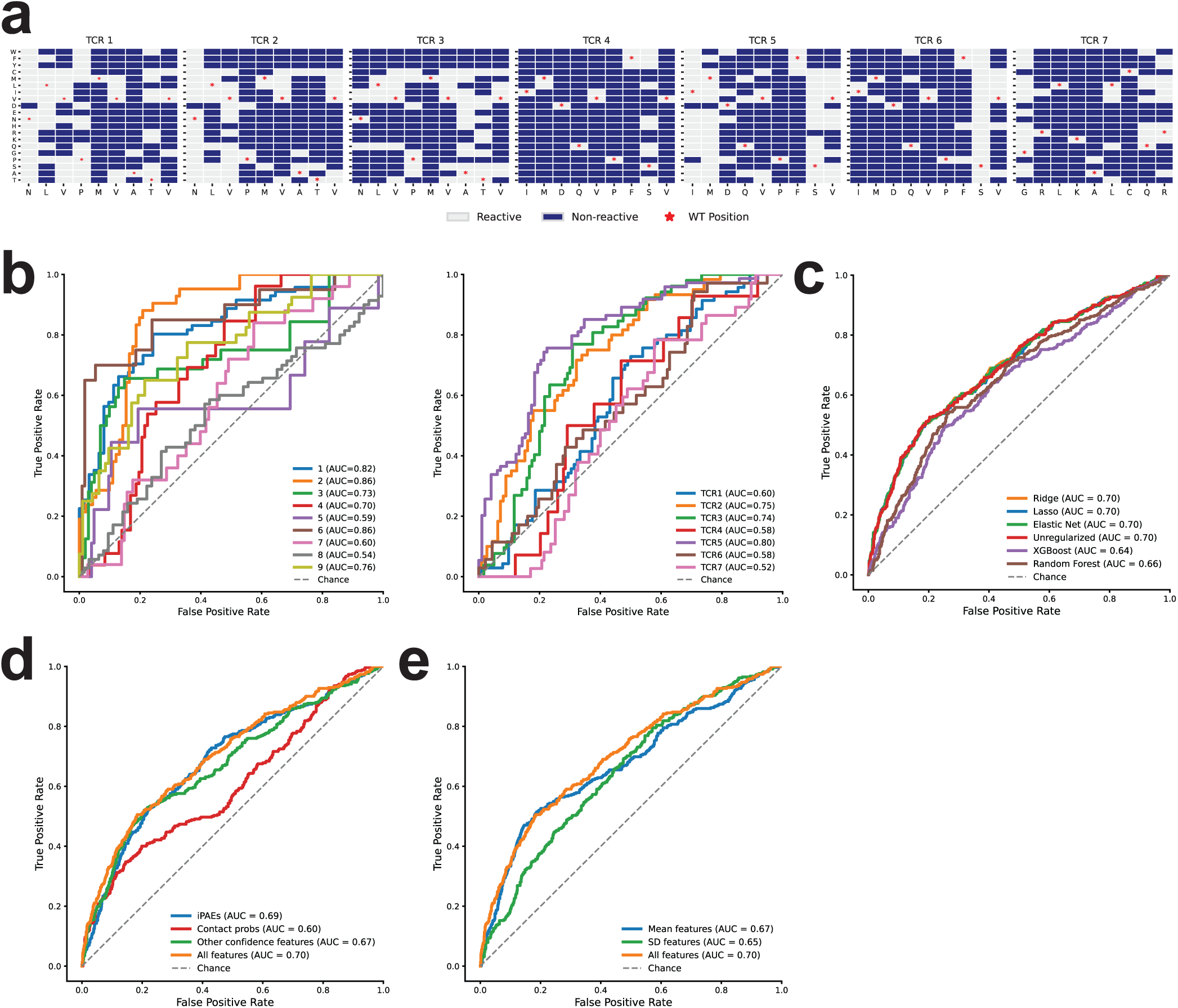
Detailed performance on the cross-reactivity dataset. **a,** Full cross-reactivity dataset with T cell reactivity to mutated peptides estimated by EC_50_ from T cell activation experiments. Reactivity is set according to the EC_50_ threshold as described in the original publication. **b,** enFoldX-Ridge model performance on cross-reactivity dataset for “Leave-One-Mutated-Position out” (**left**) and “Leave-One-TCR out” (**center**). **c,** Comparison of performance for alternative enFoldX machine learning classifiers. **d,** Comparison of performance for enFoldX-Ridge models using a subset of feature types versus including all feature types (orange)**. e,** Comparison of performance for enFoldX-Ridge models using only feature means (blue), only feature standard deviations (green), versus both feature means and standard deviations (orange) across the ensemble.

**Extended Data Fig. 7.**
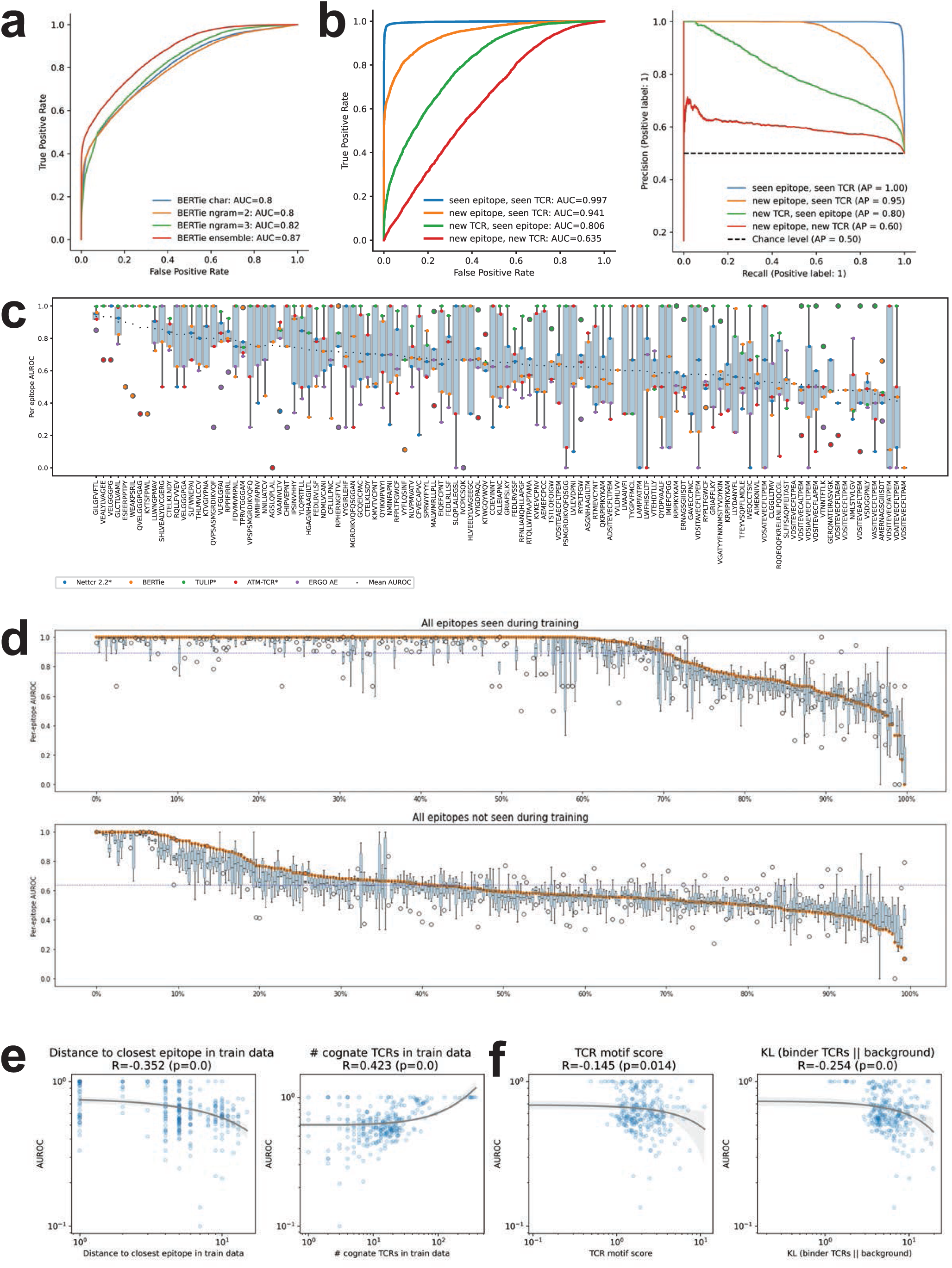
Additional analysis of sequence-based prediction model, BERTie. **a,** Performance for different BERTie encoding schemes on held-out test data. ROC curves for BERTie models encoding sequences using single amino acids (blue), 2-mers (orange), and 3-mers (green) show slightly different performance, and the ensemble of models (red) outperforms each individual sequence encoding approach. **b,** BERTie model performance on held-out test data by test scenario, as measured by ROC and precision-recall curves. AP: average precision. **c,** Per-epitope AUROC scores for BERTie and other benchmarked sequence models on the IEDB test set (epitopes with at least 2 cognate TCRs) shows lack of consistency in predictions among sequence models, including models trained on the IEDB data (indicated with *). **d,** BERTie per-epitope AUROC score is consistent among the 5 cross-validation folds for seen (**top**) and unseen (**bottom**) epitopes. Each dot is the per-epitope AUROC score of a BERTie model trained on one of the five cross-validation splits with the mean AUROC score shown in orange. Only epitopes with at least two cognate TCRs are shown. **e,** Per-epitope AUROC score for new epitopes is negatively correlated with Levenshtein distance to the most similar epitope in the training data (**left**) and positively correlated with the number of binder TCRs for each epitope included in the training data (**right**). **f,** Per-epitope AUROC score for seen epitopes is correlated with the TCR motif score that detects a consistent motif in binder TCRs in training data for this epitope (**left**) and the KL divergence between distribution of amino acids at each position in binder TCRs compared to a background, measuring how different the binder TCRs are from the rest of TCRs in training data (**right**).

**Extended Data Fig. 8.**
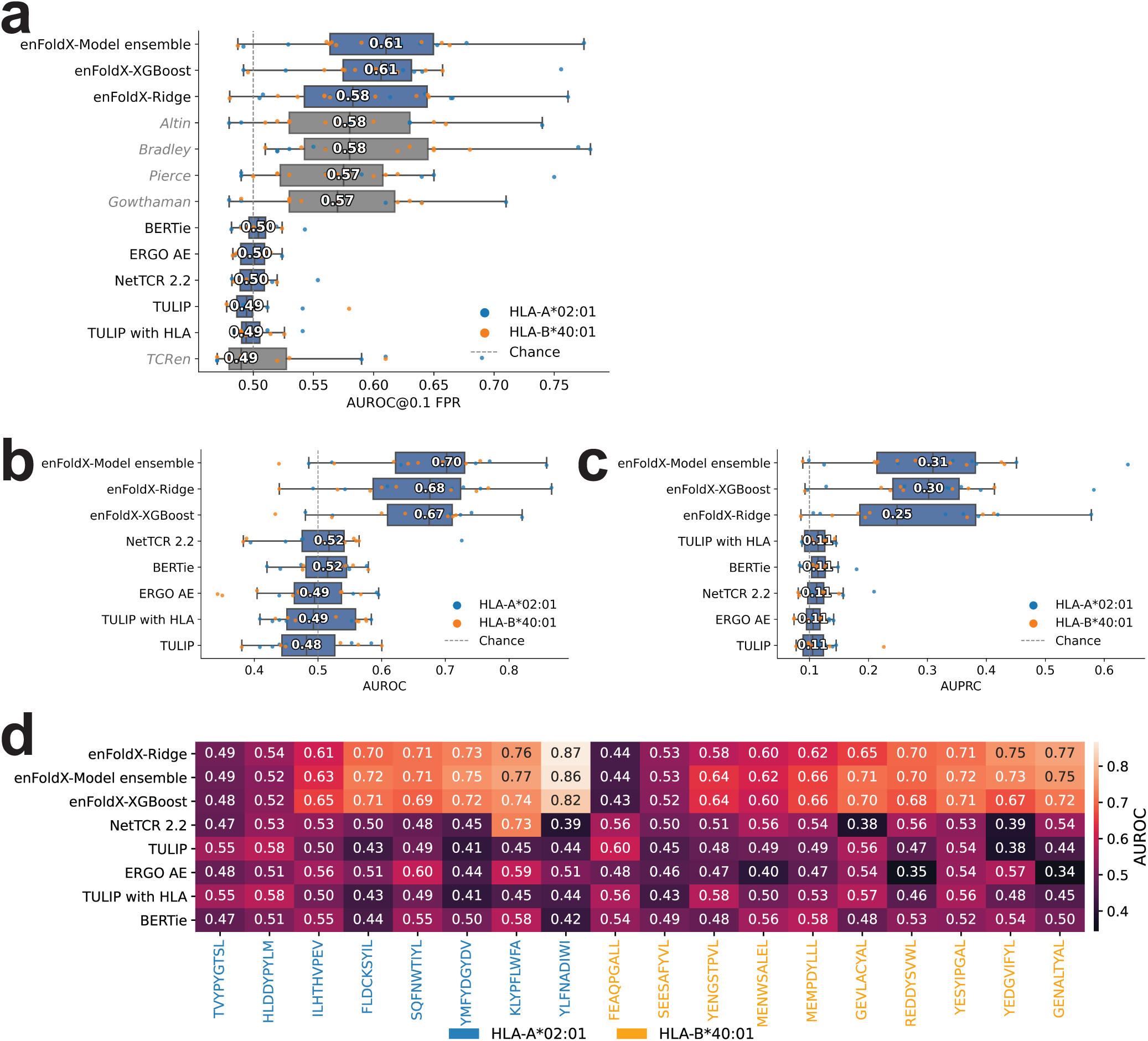
**Detailed benchmarking on the IMMREP25 test data. a-c**, Performance comparison of enFoldX and alternative methods on the IMMREP25 dataset. Methods and results reported by IMMREP25 are marked in gray. Data is shown as the AUROC@0.1 FPR (**a**), AUROC (**b**), or AUPRC (**c**) across epitopes. **d,** Per-epitope performance comparison of enFoldX and alternative methods by AUROC on the IMMREP25 dataset. Epitopes are grouped by the presenting HLA allele.

**Extended Data Fig. 9.**
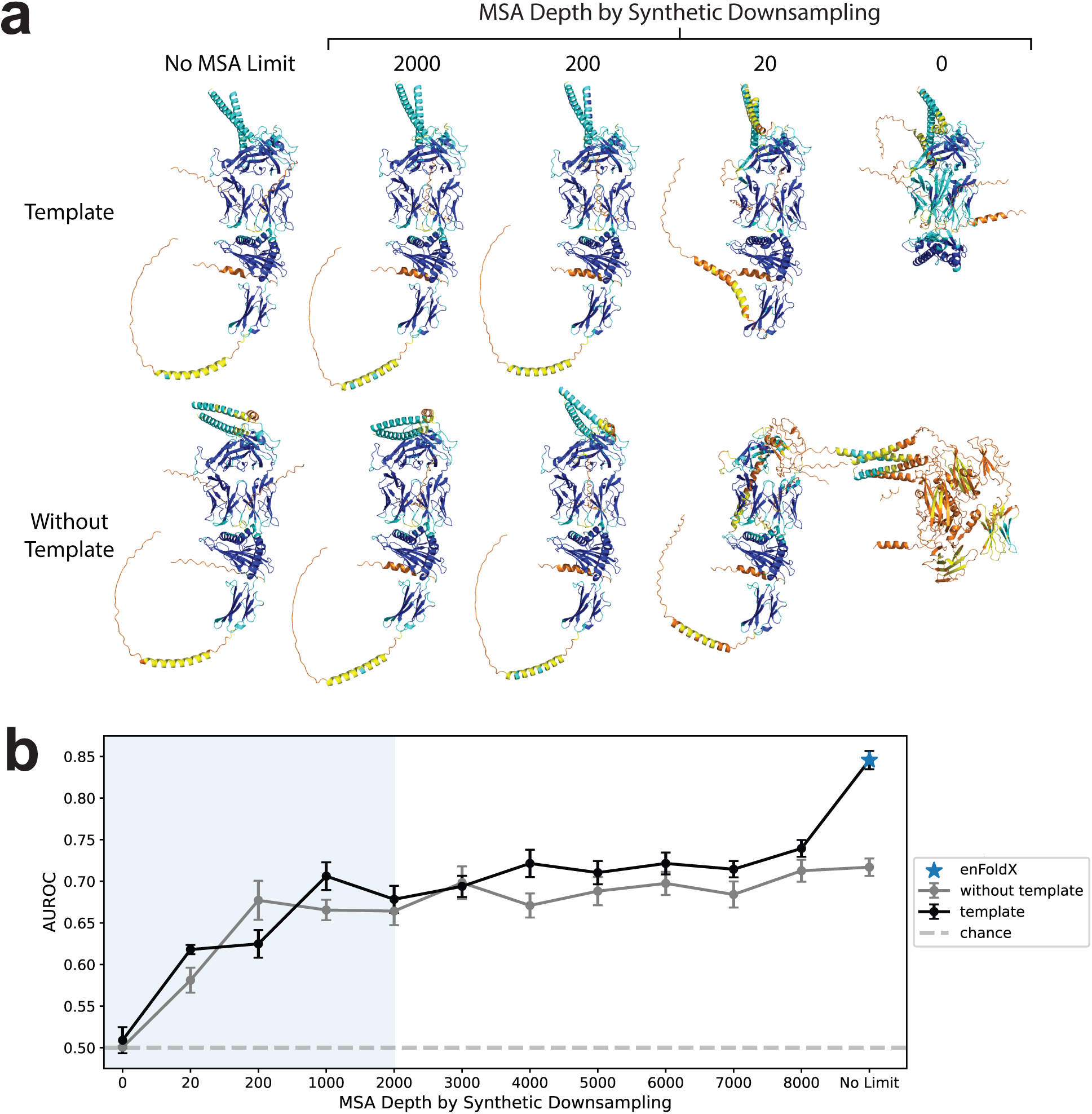
Synthetic MSA downsampling. **a**, Representative structures for TCR1 with the WT NLV peptide presented by HLA-A*02:01 from the cross-reactivity data with subsampled MSA with or without AF3 templates. **b,** Cross-validation performance on the cross-reactivity data with training using the default enFoldX pipeline (folded structures with no limit on MSA depth or templates) and testing on datasets with varying levels of downsampled MSA depth, including (black) or excluding (gray) templates. Performance at each depth is shown as AUROC +/- SEM over the folds.

**Extended Data Fig. 10.**
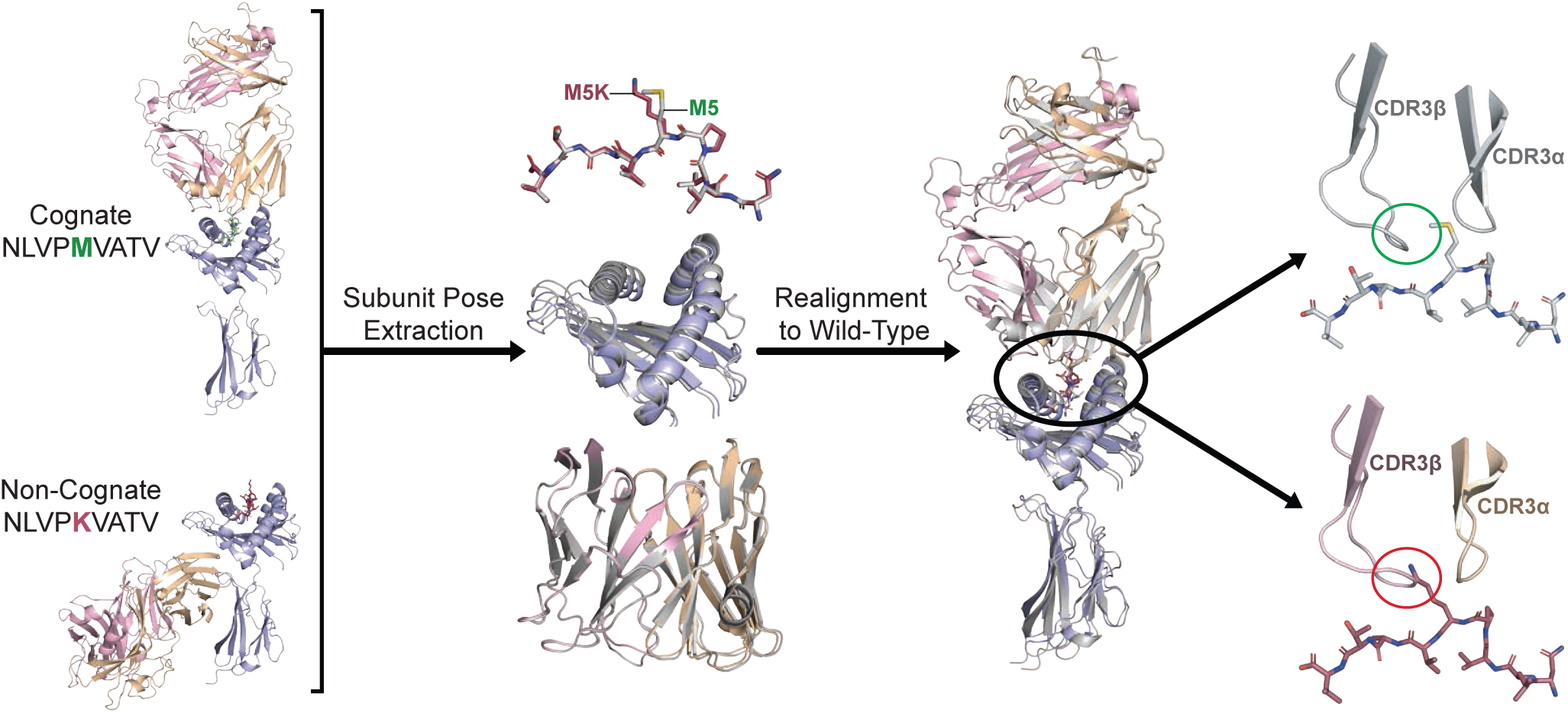
Chain re-alignment from misdocked hallucination to cognate structural prediction indicates TCR-peptide steric clash. Representative structures of TCR 5-2 with cognate WT NLV and non-cognate mutant M5K peptides from PTEAM dataset (**left**). Peptide, MHC, and TCR poses were extracted from the misdocked non-cognate structure and re-aligned to their corresponding cognate pose (**center left**) and to the overall cognate complex (**center right**). Unlike the wild-type positioning, the re-aligned pose for the M5K non-cognate pair indicates a severe steric clash between the CDR3β and the peptide, potentially explaining the physiologically impossible AF3-predicted misdocked TCR positioning.

